# Do we predict upcoming speech content in naturalistic environments?

**DOI:** 10.1101/366427

**Authors:** Evelien Heyselaar, David Peeters, Peter Hagoort

## Abstract

The ability to predict upcoming actions is a characteristic hallmark of cognition and therefore not surprisingly a central topic in cognitive science. It remains unclear, however, whether the predictive behaviour commonly observed in strictly controlled lab environments generalizes to rich, everyday settings. In four virtual reality experiments, we tested whether a well-established marker of linguistic prediction (i.e. anticipatory eye movements as observed in the visual world paradigm) replicated when increasing the naturalness of the paradigm by means of i) immersing participants in naturalistic everyday scenes, ii) increasing the number of distractor objects present, iii) manipulating the location of referents in central versus peripheral vision, and iv) modifying the proportion of predictable noun-referents in the experiment. Robust anticipatory eye movements were observed, even in the presence of 10 objects (hereby testing working memory) and when only 25% of all sentences contained a visually present referent (hereby testing error-based learning). The anticipatory effect disappeared, however, when referents were placed in peripheral vision. Together, our findings suggest that working memory may play an important role in predictive processing in everyday communication, but only in contexts where upcoming referents have been explicitly attended to prior to encountering the spoken referential act. Methodologically, our study confirms that ecological validity and experimental control may go hand in hand in future studies of human predictive behaviour.

## 1. Introduction

In the last few decades, there has been an increased interest in the role of prediction in language comprehension. The idea that people predict (i.e., context-based pre-activation of upcoming linguistic input) was deemed controversial at first: With Humboldt’s characterization of language as a system that “makes infinite use of finite means” (1829), there would be an infinite list of possibilities, rendering it inefficient to predict upcoming information (Jackendoff, 2002). However, current theories of language comprehension have embraced linguistic prediction as the main reason why language processing tends to be so effortless, accurate, and efficient (see Clark, 2013; Friston, 2010 for an overview). Current theories of prediction involve the creation of an internally generated model of anticipated upcoming information, similar to the efference copy proposed to drive prediction in the motor movement field (see Wolpert & Flanagan, 2001). The actually encountered linguistic information is then compared against this forward model of anticipated linguistic information (Pickering & Garrod, 2007) and any prediction error is used as a learning mechanism that influences future predictions (Dell & Chang, 2014). The theories are mostly based on data collected via EEG and the visual world paradigm, the latter of which we will focus on in this study.

The visual world paradigm (VWP) is based on the phenomenon that when participants are presented with spoken language while viewing a visual scene, their eye movements are very closely synchronized to a range of different linguistic events in the speech stream (Cooper, 1974). Altmann and Kamide (1999) exploited this behaviour to illustrate that participants anticipate upcoming linguistic information during language comprehension. In their seminal study, participants were presented with a visual scene depicting, for example, a boy, a cake, and some toys. While participants heard sentences such as “the boy will move the cake” or “the boy will eat the cake", the authors observed that participants would fixate on the cake significantly earlier after hearing the verb form “eat” (but before “cake” was uttered) compared to after hearing the verb form “move". The VWP has since been proven as an excellent method to provide direct evidence of what type of information is anticipated (Coco & Keller, 2015; Hintz, Meyer, & Huettig, 2017; Kamide, Altmann, & Haywood, 2003; Knoeferle & Crocker, 2006, inter alia). Other commonly used methods, such as EEG, have also provided ample evidence in favour of linguistic prediction, although it is debatable whether several observed effects truly reflect a measurement of prediction or rather reflect integration of encountered input with the preceding context instead (Kochari & Flecken, 2018; Kutas, DeLong, & Smith, 2011; Nieuwland et al., 2018; van den Brink, Brown, & Hagoort, 2001). The elegance of the VWP is its ability to measure the direct interaction of language and the visual world, not surprisingly making it a commonly used paradigm.

One principle question that has received very little attention is *when*, i.e. in which everyday contexts, we actually predict (Huettig, 2015). Although ample VWP evidence has shown that we are able to predict, whether we do so in naturalistic settings, and whether we do it all the time, are open questions that have been difficult to answer with traditional methods. The aim of the current study is therefore to use the VWP to provide empirical evidence to answer this *when* question, and, while doing so, we also test for characteristics of prediction that have, as of yet, received little or no empirical support. We will initially re-design elements of the VWP in order to make it more naturalistic (e.g. by introducing a source for the spoken sentences, placing objects in thematic settings, etc., see below). This new set-up will allow us to manipulate not only the realism of the scenes, but also manipulate other aspects, such as frequency of predictable sentences and number of objects in the scene. We can then use this new set-up to provide evidence for the use of working memory (by increasing the number of distractor objects present and comparing VWP performance to the participant’s working memory capacity) as well as error-based learning in prediction; two elements the current theories of prediction are built upon, but for which little empirical evidence exists. For the remainder of the introduction we will provide further background and details for each of our aims.

### 1.1 Experiment 1 - Improving the VWP

Although the VWP is commonly considered as a relatively naturalistic way to measure the interaction between language and visual attention, it is not without its limitations. Participants are seated in front of a computer screen and presented with 2D objects, which, in the majority of experiments, are four simple line drawings presented 2×2 on a white background. Indeed, if tested with larger arrays (3×3 or 4×4), no anticipatory eye-movement is observed (Sorensen & Bailey, 2007). The sentences are always played from a disembodied voice and if human characters are present in the scene, great effort is made to control for the positioning and direction of gaze of the character such that it does not show favour to any particular object (Coco, Keller, & Malcolm, 2016; Eichert, Peeters, & Hagoort, 2017). Additionally, participants are played only one sentence per scene/array, which often makes the goal of the study (detecting saccades made to a mentioned object) salient to the experimental participant.

These limitations in terms of ecological validity call into question whether the behaviour exhibited in the lab is reflecting natural behaviour or is perhaps induced by the rigorously controlled experimental environment. In short, what previous studies show is what listeners *can* do, not necessarily what they actually do in naturalistic contexts. By definition, this therefore also calls into question the generalizability of the theories developed about linguistic prediction that have been based on results from the VWP. The aim of the current experiment is therefore to increase the ecological validity of the VWP to determine whether anticipatory eye-movements are indeed a natural phenomenon or an experimental artifact.

We conducted this study in Virtual Reality (VR) to ensure participants feel immersed in the environment while retaining the required levels of experimental control for reliable data collection. By immersing participants in a 3D world instead of presenting 2D cartoon images on a computer screen, the immediate goal of the study becomes less obvious to the participants, and therefore encourages more naturalistic behaviour. Previous studies have built VR versions of common psycholinguistic tasks and have shown comparable behaviour with the traditional version (Heyselaar, Hagoort, & Segaert, 2015; Peeters & Dijkstra, 2017; Tromp, Peeters, Meyer, & Hagoort, 2017). Recently, Eichert and colleagues (2017) moreover showed robust anticipatory eye-movement behaviour in a VR version the VWP task, suggesting that using 3D versus 2D objects in itself does not change participants’ anticipatory eye-movement behaviour. The current study will go several steps further in making use of the unique affordances of VR and increase the naturalness of the VWP in ways that are hard or impossible to imagine in traditional versions.

The backbone of the experimental set-up is the immersion of participants in realistic everyday scenes such as a living room, an office, a neighbourhood, etc. As a first step towards mimicking real-world communication, participants will be taken on a tour by a virtual agent, who will also deliver the critical sentences. To ensure that participants remain unaware of the goal of the study, each scene will contain four critical sentences (compared to one sentence per scene as done in traditional versions). Therefore, to minimize any benefits of guessing, we have increased the number of objects from the traditional 4 to 6. Experiment 1 tests this initial set-up to ensure the changes we have made do not already discourage anticipatory eye-movement behaviour. The remainder of the experiments in this study manipulate aspects of this set-up, such as the number of distractor objects or the predictability of the sentences, to build towards a more accurate reflection of real-world situations. Whether participants predict in these situations has already been mapped out in theories of linguistic prediction: Working memory capacity should dictate how participants perform with increased distractors, and error-based learning should dictate how participants would perform if the predictability of the sentences were manipulated. Therefore, our study not only addresses *when* we predict, but also provides empirical evidence whether working memory and error-based learning do in fact play a role in linguistic prediction in natural, everyday settings.

### 1.2 Experiments 2 and 3 - The Role of Working Memory in Prediction

In Experiment 2 we explore the effect of increasing visual complexity by increasing the number of objects presented in each scene. Previous studies are conflicting on whether increased visual complexity affects anticipatory eye-movement behaviour. As discussed above, Sorensen and Bailey (2007) observed a lack of effect when presenting participants with 9 or more items. In contrast, studies using complex, photographic scenes have still shown language-driven eye-movements (Andersson, Ferreira, & Henderson, 2011; Coco & Keller, 2015; Coco et al., 2016). Additionally, increasing the number of objects present in the scene would also tax working memory, a process that has been theorized to play an important role in prediction.

There is a growing consensus for a role of working memory in prediction (see for review Huettig, Olivers, & Hartsuiker, 2011). In the VWP, anticipatory looks to the referent object can occur as early as 200ms after the verb is heard, a timeframe that suggests that participants already had the potential objects activated to some extent. Huettig and colleagues propose that objects in the display are first encoded in a visuospatial type of working memory (cf. Baddeley, 1998; Cavanagh & Alvarez, 2005; Pylyshyn, 1989), which triggers perceptual hypotheses in long-term memory. These hypotheses then trigger a cascade of activations of associated semantic and phonological codes, all within a few hundreds of milliseconds (cf. Huettig & McQueen, 2007). This results in a nexus of associated knowledge, which is bound to an object’s location within working memory. Hence object selection and planning a saccade to the location of said object is faster due to the already activated representations within working memory. Indeed, a recent study showed a correlation between a working memory construct and predictive looks towards 4 objects (Huettig & Janse, 2016).

However, working memory has a limited capacity: The capacity of visual working memory has been shown to be between 3 and 4 items (Luck & Vogel, 1997; Vogel & Machizawa, 2004). As current VWP experiments commonly limit the number of potential objects to 4, the capacity limit is barely crossed. If working memory indeed plays a role in prediction, one would assume that by increasing the number of potential object referents in a visual scene, the anticipatory eye movement behaviour will decrease as participants can no longer accurately maintain all the objects’ representations online. Additionally, this behaviour should be modulated by the participant’s individual working memory capacity. As such, we will additionally measure each participant’s visual working memory capacity, to determine whether participants with lower working memory capacity indeed exhibit less anticipatory eye movements compared to participants with higher working memory capacity.

If the VWP is indeed an ecologically valid methodology to study the interaction of language and the visual world, then one would predict that anticipatory eye movements also occur in the real world. Therefore, by definition, only one of these two alternatives can be true. If prediction involves working memory then prediction-ability should decrease once the working memory limit is exceeded. Hence, outside the laboratory, we would expect very little anticipatory eye movements as we are rarely in an environment with four or less objects. On the other hand, if the VWP does measure ecologically valid prediction behaviour, then participants should also exhibit anticipatory eye movements in rich, complex scenes with more than 4 objects, which cannot be possible if fully supported by working memory.

A key requirement for involving working memory is that the objects must already be in working memory in order to be selected. Therefore, in Experiment 3, we moved referent objects into the visual periphery such that they are not automatically encoded by the participant. If another process is responsible for anticipatory eye movement behaviour, such as perhaps a visual search for a contextually relevant referent upon hearing a potentially restrictive verb, then we predict to still see fixations on the referent object in this experiment.

### 1.3 Experiment 4 - Error-based Learning

Increased realism is not limited to visual complexity. As displacement is an important and common feature of present day human communication (Hockett, 1960), not every sentence in a conversation necessarily refers to an object in the interlocutor’s immediate environment. Therefore, in Experiment 4, we included filler sentences that referred to objects not present in the scene. The distribution was such that per scene, only 50% of the sentences referred to objects present, and only 25% of the total sentences utilized verbs that allowed the noun to be predicted on the basis of the visual context. This manipulation therefore also tests whether participants would adapt their prediction behaviour when they realize that the majority of the referential nouns cannot be predicted and therefore it would be inefficient to try.

Although there are many different proposed mechanisms for prediction (G. T. Altmann & Mirkovic, 2009; Chang, Dell, & Bock, 2006; Dell & Chang, 2014; Kahneman, 2011; Kuperberg, 2007; Pickering & Garrod, 2007), the majority propose that prediction makes use of previous experience. Events tend to reoccur and show regularities and therefore are likely to be an important organizing principle of past experience. As described in Dell & Chang (2014, p. 4): “the central component of the model tries to predict the next heard word from the word that preceded it and a representation of prior linguistic context. It then compares the predicted next word with the actual next word. The resulting prediction error is used to change the model’s internal representations, thus enabling the model to acquire the knowledge that helps it make these predictions.”

Therefore, if past experience does play a role in predicting upcoming information, then we would assume that participants would show less anticipatory behaviour in Experiment 4, as any predictions they attempt to make regarding an upcoming noun-referent on the basis of the immediate visual environment will only be relevant and correct 25% of the time.

### 1.4 Overall aim

In sum the VWP is an important methodology used to investigate linguistic prediction. Although it aims to measure ecologically-relevant behaviour, it comes with several limitations that may have encouraged behaviour that the average person may not produce in the real world. Therefore, in this study we create a more realistic VWP by placing participants in 3D worlds with thematic objects and a narrator. By increasing the number of objects and manipulating how often participants hear a sentence with a predictable noun-referent, we not only measure anticipatory eye-movement in real-world contexts, but we are also able to empirically test whether elements such as working memory and error-based learning do indeed play an important role in linguistic prediction in everyday settings.

## 2. Experiment 1: Improving the Visual World Paradigm

### 2.1 Materials and Methods

This study and experiments 1, 2, and 4 were pre-registered via Open Science Framework and can be found under the title: “Language-driven anticipatory eye-movements in naturalistic settings.” Experiment 3 and Overall Results were not pre-registered and therefore fully exploratory.

#### 2.1.1 Participants

Twenty native speakers of Dutch (13 female, M_age_: 22.8 years, SD_age_: 3.50 years) were recruited from the Max Planck Institute for Psycholinguistics database. The data of 24 participants was recorded, but one participant was discarded due to insufficient accuracy of the eye-tracking data and three stated during the debrief stage that they did not understand the virtual agent properly (clarity rating < 3 out of 5). The participants gave written informed consent prior to the experiment and were monetarily compensated for their participation.

#### 2.1.2 Materials

##### 2.1.2.1 Virtual agent

The virtual agent was adapted from a stock virtual agent produced by WorldViz (2016; “casual03_f_highpoly”). The virtual agent’s appearance suggested that she was a Caucasian female in her mid-twenties, which matched the age and ethnicity of the native Dutch speaker who recorded her speech. All the virtual agent’s speech was pre-recorded.

##### 2.1.2.2 Scenes

Eight scenes were designed to represent places in the virtual agent’s life (her office, her neighbourhood, her living room, etc.; see Appendix for a full list of scenes). The scenes were designed to appear as realistic as possible (Figure 1) but initially without any objects. Scenes that came with furniture (such as the table in Figure 1) were designed such that these items would not be predictable given the sentences. Objects were then placed in realistic locations in each scene. For example, the car in the neighbourhood scene was placed in the driveway, the tree was placed on the grassy lawn, and the basketballs were placed on the sidewalk. The aim was to place the objects in such a way that they were not overly salient, however, objects always appeared in the middle screen so that participants did not have to search for them.

**Figure 1.**
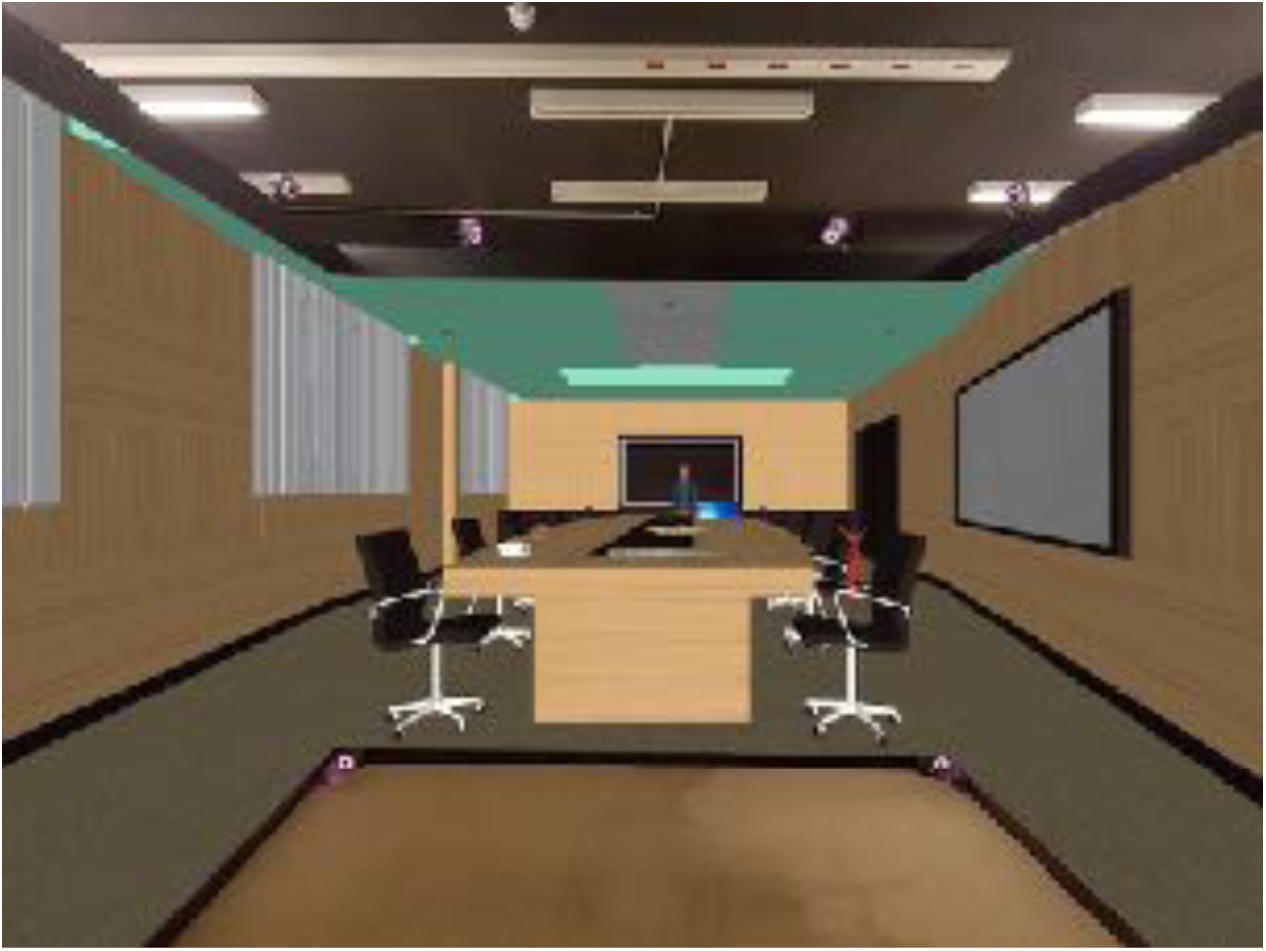
Example scene. The figure illustrates the office scene, one of eight scenes used in this study. Participants, wearing 3D-glasses, stood in front of the virtual desk during the task, and were allowed to move their heads to view the entire scene. The location of the two screens to the left and right gives the participants a feeling of being immersed in the scene. The virtual agent would always be present on the middle screen, to ensure participants were able to locate her easily and would feel addressed when she spoke to them. The 6 objects present (cup of coffee, muffin, newspaper, pizza, laptop, and tie) are placed in naturalistic locations and scaled to realistic proportions relative to the room. Participants heard four sentences while viewing this scene, two of which were restrictive (e.g., *after work, sometimes someone drinks a cup of coffee)* and two of which were unrestrictive (e.g., *if it’s busy, we share a laptop)*. In this case, a pre-test indicated that participants found the cup of coffee the only drinkable object in the scene, yet they found find the cup of coffee, laptop, pizza, and muffin shareable.

The virtual agent appeared in each scene in the middle screen such that participants would feel addressed when she spoke to them.

##### 2.1.2.3 Objects and sentences

Thirty-six sentence pairs were created, of which one sentence was *restrictive* (the verb imposed constraints on its arguments such that only one of the visually presented objects was a plausible completion of the sentence) and one was *unrestrictive* (no such constraints were imposed; the sentence could be completed with at least three of the objects present in the scene). The sentence pairs therefore only differed in their verb. For example, *na werktijd <u>drinkt</u> soms iemand een kopje koffie* ("after work, sometimes someone <u>drinks</u> a cup of coffee”) versus *na werktijd <u>haalt</u> soms iemand een kopje koffie* ("after work, sometimes someone <u>gets</u> a cup of coffee"; see Appendix for a full list of sentences and their English translations).

All the objects present in the experiment were selected from a standardized database of 3D objects (Peeters, 2017) to ensure that all objects were easily identifiable. The experiment contained eight scenes. Each scene included four sentences; six objects were present in each scene. This ensured that even with the fourth sentence, participants still had three objects that had not yet been mentioned, and therefore they could not accurately guess the target object for the final sentence.

The verbs were chosen such that their word length and frequency were not significantly different between conditions (length: Mann-Whitney *U* = 416, *p* = .189; frequency: Mann-Whitney *U* = 475, *p* = .619).

Thirty-eight participants (who were not invited for the main experiment) completed an online Cloze-like task to ensure that the target object was the most likely completion for the restrictive sentences (M: 92.67%, SD: 18.98%) compared to the unrestrictive sentences (M: 19.51%, SD: 21.21%). Participants were given the incomplete sentence and asked to choose the most likely completion from a list of the objects in the scene.

Sentences were recorded in a sound-proof booth, sampling at 44.1 kHz (stereo, 16 bin sampling resolution). All files were equalized for maximal amplitude. Sentences were annotated using Praat (Boersma & Wennink, 2009) by placing digital markers at onsets and offsets of critical words: Verb onset, verb offset, noun onset, noun offset, and end of sentence. The mean duration of the sentences was 2,474ms. During recording of the sentences, we ensured an average of 571ms (SD: 116ms) between the end of the verb and the start of the noun (time of interest [TOI]), as previous research has shown that at least 500ms is necessary to successfully allow prediction effects (Salthouse, McGuthry, & Hambrick, 1999). We observed no significant difference in the length of the TOI between the two conditions (*t*(62) = −0.51, *p* = .612).

#### 2.1.3 Apparatus

##### 2.1.3.1 CAVE system

The experiment was run in a CAVE Virtual Reality set-up (see Figure 1), the layout of which has been described before in detail in Eichert and colleagues (2017). The CAVE system consisted of three screens (255 cm x 330 cm, VISCON GmbH, Neukirchen-Vluyn, Germany) that were arranged at right angles. Two projectors (F50, Barco N.V., Kortrijk, Belgium) illuminated each screen indirectly through a mirror behind the screen. The two projectors showed two vertically displaced images which were overlapping in the middle of the screen. Thus, the complete display on each screen was only visible as combined overlay of the two projections. For optical tracking, infrared motion capture cameras (Bonita 10, Vicon Motion Systems Ltd, UK) and Tracker 3 software (Vicon Motion Systems Ltd, UK) were used. The experiment was programmed and run using 3D application software (Vizard, Floating Client 5.4, WorldViz LLC, Santa Barbara, CA), which makes use of the programming language Python. Sound was presented through two speakers (Logitech, US) that were located at the bottom edges of the middle screen. For a schematic of the CAVE set-up, see Eichert and colleagues’ (2017) Figure 4.

##### 2.1.3.2 Eye-tracking

Eye-tracking was performed using special glasses (SMI Eye-Tracking Glasses 2 Wireless, SensoMotoric Instruments GmbH, Teltow, Germany) that combine the recording of eye gaze with the 3D presentation of VR. The recording interface used was a tablet that was connected to the glasses by cable. The recorder communicated with the externally controlled tracking system via a wireless local area network, which enabled live data streaming.

The glasses were equipped with a camera for binocular 60 Hz recordings and automatic parallax compensation. The shutter-device and the recording interface were placed in a shoulder bag worn by the participants. This enabled the participants to move freely through the CAVE if they so chose. In reality, the participants stayed standing in the center of the room, roughly 180cm away from the central screen. Gaze tracking accuracy was estimated by the manufacturer to be 0.5° over all distances. We found the latency of the eye-tracking signal to be 60ms ± 10ms. This latency was corrected for in the statistical analyses (see below).

By combining eye-tracking and optical head-tracking, we were able to identify the exact location of participants’ eye gaze in three spatial dimensions, allowing them to move their heads during the experiment. Optical head-tracking was accomplished by placing light reflectors on both sides of the glasses. Three spherical reflectors were connected on a plastic rack and two of such racks with a mirrored version of the given geometry were manually attached to both sides of the glasses using magnetic force. The reflectors functioned as passive markers which were detected by the infrared tracking system in the CAVE. The tracking system was trained to the specific geometric structure of the three markers and detected the position of the glasses with an accuracy of 0.5mm.

##### 2.1.3.3 Regions of Interest

In order to determine target fixations, we defined individual 3D regions of interest (ROIs) around each object in the virtual environment. The *x* (width) and *y* (height) dimensions of the ROI were adopted from the frontal plane of the object’s individual bounding box, facing the participant. We adjusted the size of this plane to ensure a minimal size of the ROI. The minimal width was set to 0.8 and the minimal height to 0.5. For the presented layout of objects, the adjusted *x* and *y* dimensions were sufficient to characterize the ROIs. Despite the 3D view, the plane covered the whole object sufficiently to capture all fixations. The *z* dimension (depth) of the ROI was therefore set to a relatively small value of 0.1. An increased *z* value of the ROIs would not have been more informative about the gaze behaviour, but would have led to overlapping ROIs in some cases. The eye-tracking software automatically detected when the eye gaze was directed to one of the ROIs and coded the information online in the data stream. Some previous studies have used contours of the objects to define ROIs, but rectangles have been shown to produce qualitatively similar results (G. T. M. Altmann, 2011). In addition to the six objects in each scene, an ROI was also coded for the virtual agent.

#### 2.1.4 Design and Procedure

Participants were instructed to stand in the middle of the CAVE system, roughly 180cm away from the middle screen. They put on the VR glasses, which were softly fastened using a strap on their head to ensure stability. Prior to the start of the experiment, two calibration steps were performed. For the first calibration step, calibration was done using the SMI software ‘ One-step Calibration’ program. The second calibration step is as described by Eichert et al. (2017): Participants were asked to look at three displayed spheres successively. The experimenter selected the corresponding sphere. The computer software computed a single dimensionless error measure of the eye-tracker combining the deviance in all three coordinates. The computer-based calibration was repeated until a minimal error value (< 4) and thus maximal accuracy was reached.

Deviance was checked during the break and re-calibrated using the three-spheres procedure if the error value was greater than 4. This was only necessary for one participant.

Prior to the start of the experiment, participants were informed that they would be given a tour of a virtual agent’s life and that the goal of the experiment was to form an opinion of the virtual agent. After the virtual reality portion, they were told they would be given a questionnaire asking for their opinion of the virtual agent. This insured that the participants paid attention to the virtual agent and drew potential attention away from the objects. During the debrief stage, none of the participants had guessed at the purpose of the experiment, although one participant thought that they had to memorize which objects were present in each scene.

Participants were presented with two experimental blocks of four scenes (32 sentences) each. The first block contained the office, forest, café, and canteen scene; the second block contained the living room, bathroom, attic, and neighbourhood scene (see Appendix). All scenes were randomized within each block for each participant, although the living room scene was always the first scene presented in the second block for all participants (see below). Each scene had a preview time of 1s before the virtual agent gave a short introduction (M: 2.02s), after which there was a 2.5s wait time before the first sentence was played. This gave participants an average of 4.5s preview time of each scene. For the living room scene, the virtual agent’s introductory text was “welcome to my house” and hence it was always the first scene of that block. The task took around 7 minutes to complete.

We created two lists of 32 restrictive sentences and 32 unrestrictive sentences taken from each sentence pair. No list contained both the restrictive and unrestrictive versions of the same sentence pair. Participants were assigned to a list based on their participant number (odd participants were assigned to list 1; even participants were assigned to list 2). Sentences were presented randomly within each scene for each participant. As the last sentence presented in each scene meant that the participants had had a maximal viewing time of the scene and its objects compared to the first sentence presented, by randomizing the sentences, this balanced out any beneficial effects across the experiment.

Participants were given a self-timed break after the fourth scene. During this time participant’s calibration was checked and re-calibrated if necessary. Calibration was also checked at the end of the experiment. After the experiment, participants were given a debrief questionnaire in which they were asked to rate the clarity of the virtual agent’s speech as well as indicate which objects they heard the virtual agent refer to. This list contained all the objects present in the experiment, of which only 66.67% were actually named by the virtual agent. Accuracy on this questionnaire was taken as an indication of how well the participants paid attention to what the virtual agent was saying.

#### 2.1.5 Statistical Analyses

Data was acquired at a sampling frequency of 60 Hz. We corrected for the 60ms latency shift caused by the eye-tracking system by time-locking the data to 60ms (~4 frames) after each sentence onset. A fixation was defined as a look to the same ROI that lasted at least 100ms. This correction on the experimental data led to an exclusion of 6.93% of all frames logged as object fixations, and 2.36% of all frames logged as virtual agent fixations. Fixation data was then aggregated into time bins of 50ms (i.e. three data frames).

We followed the steps outlined in Porretta and colleagues (2017) for analysing visual world paradigm data with general addictive mixed models (GAMM). The analysis was conducted using the *mgcv* package (version 1.8-22; Wood, 2017) and *itsadug* package (version 2.3; van Rij, Wieling, Baayen, & van Rijn, 2017) in R (version 3.4.2; R Core Development Team, 2011). As the dependent variable we entered the empirical logits of the proportion of target fixations. The model included factor smooth interactions for *Time* by *Subject*, factor smooth interactions for *Time* by *Sentence*, as well as a smooth for *Time* by *Condition* (restrictive versus unrestrictive). We included *Condition* as a parametric component, which is necessary to estimate the time curve for each level of *Condition*. We also included weighted linear regression over empirical logits as weights in the model (Barr, 2008). After fitting the model we determined an appropriate value for the AR1 parameter using the start_value_rho to account for autocorrelation in the residuals (i.e., error). We used the function plot_diff to approximate the time intervals of significant differences between conditions.

### 2.2 Results

Participants were able to accurately identify which objects the virtual agent had named and which she had not named 90.25% of the time (SD: 8.19%). Therefore we are confident that all participants listened to the virtual agent throughout the experiment.

#### 2.2.1 Inspection of the Grand Mean

To determine whether participants fixated on the target object at all during each trial, regardless of condition, we plotted the proportion of target fixations collapsed over all participants, trials, and conditions (Figure 2A). For this figure, each trial is time-locked to verb onset to give an accurate indication of eye movement behaviour in the moments after the verb is comprehended. Visual assessment of the grand mean shows a robust increase in the proportion of fixation to the target object after the noun was mentioned.

**Figure 2.**
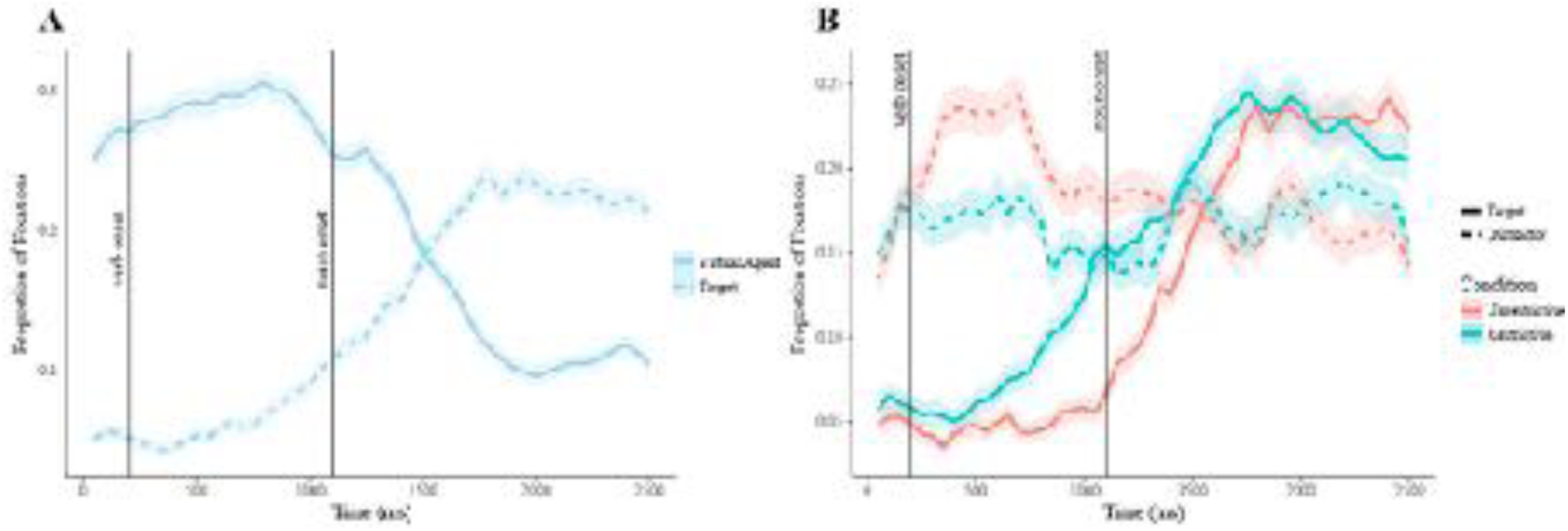
Mean proportions of fixations. **A**. To the target object and virtual agent. **B**. To the target and distractor objects shown per condition. Vertical lines indicate critical time points. The main statistical analysis was performed on the interval between verb onset and noun onset. Error clouds indicate standard error.

#### 2.2.2 Effect of Condition

For the main statistical analysis we defined a critical time window where we expected the experimental manipulation to have an effect on the proportion of target fixations. We chose the onset of the critical window as 200ms after verb onset, assuming that it takes approximately 200ms to plan and initiate a saccadic movement (Matin, Shao, & Boff, 1993). As offset of the critical time window we chose the average onset of the noun (900ms after verb onset, in line with previous studies (G. T. M. Altmann & Kamide, 1999; Eichert et al., 2017)).

We performed a generalized additive mixed model (GAMM) analysis as outlined in Porretta and colleagues (2017). The model included factor smooth interactions for *Time* by *Subject*, factor smooth interactions for *Time* by *Sentence*, as well as a smooth for *Time* by *Condition* (restrictive versus unrestrictive). We included *Condition* as a parametric component. Table 1 provides a model summary.

**Table 1.**
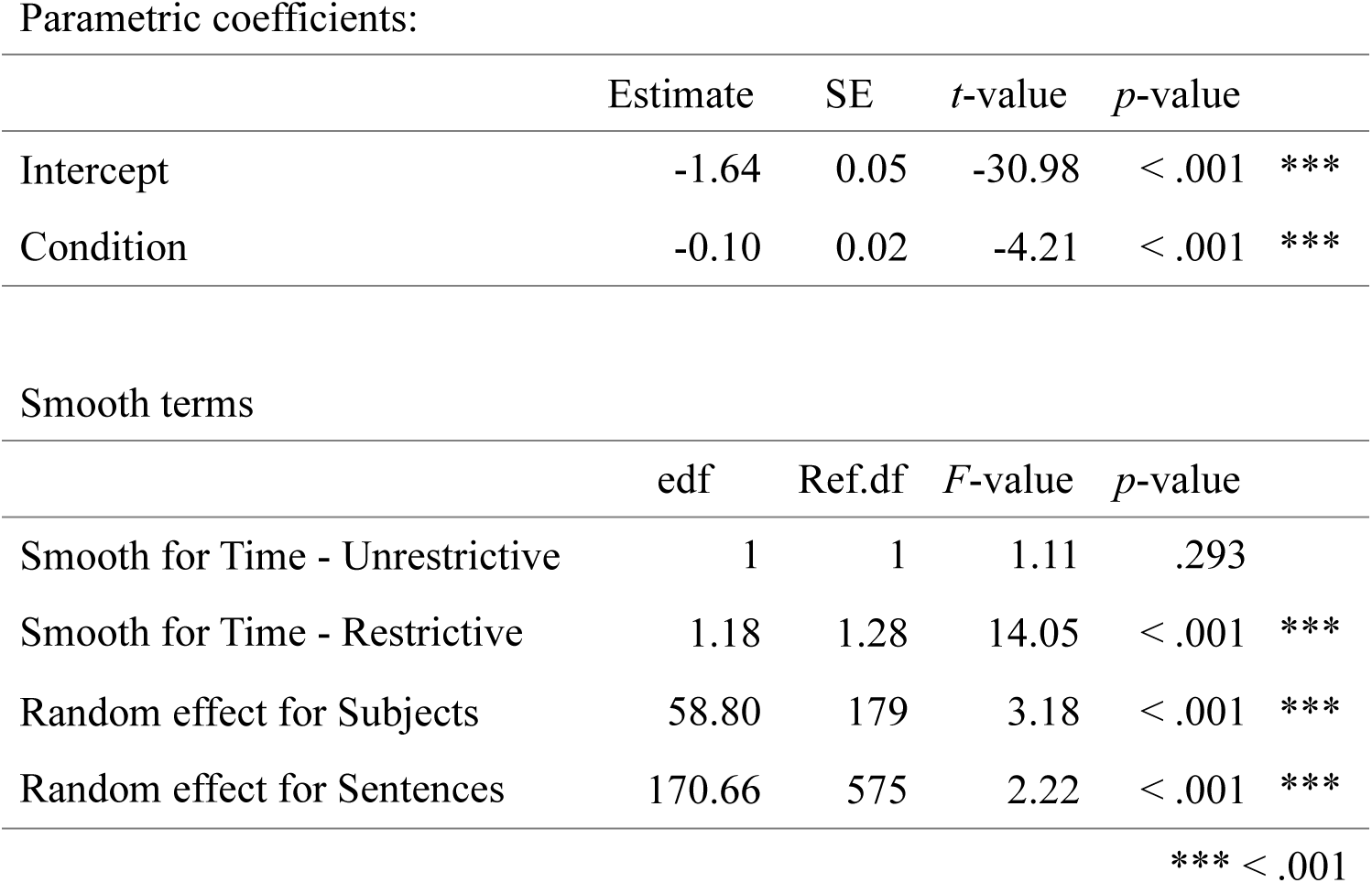
Summary of the generalized additive mixed model for changes in target fixations over time, per condition (restrictive versus unrestrictive sentences). Effective degrees of freedom (edf), reference degrees of freedom (Ref.df)

The model revealed that the curve for the restrictive condition as a function of time was significantly different from zero, whereas this was not the case for the unrestrictive condition (*p* = .293). The difference in the predicted values was significant between 398ms and 900ms after verb onset. Fixation proportions time-locked to verb onset are illustrated in Figure 2B.

We performed the same analysis on the mean distractor fixations. The model revealed the same effects, except that now there was a significant effect for the unrestrictive condition (p = .011) and not the restrictive condition (*p* = .329). The difference in the predicted values became significant between 314ms and 900ms after verb onset.

There was no effect of condition on the proportion of virtual agent fixations (p > .203).

These results are consistent with the hypothesis that participants directed their gaze towards the target object before noun onset in the restrictive condition, but not in the unrestrictive condition. Complementary to the target fixations, fixations to the distractor objects revealed that participants fixated more on distractor objects during the unrestrictive condition compared to the restrictive condition.

## 3. Experiment 2: Increased objects

Experiment 1 showed the standard anticipatory eye movement effects seen in the literature (Altmann & Kamide, 1999; Eichert et al., 2017), even in the more realistic setting our virtual reality system provided. As a next step, we enhanced the realism of our scenes by increasing the number of objects in each scene from 6 to 10. For each sentence the participants will therefore have to select from 10 potential objects within 500ms. Theories of prediction suggest a role of working memory such that participants keep potential objects ‘online’ to allow for a quick selection before the object is named (Huettig, Olivers, et al., 2011). By increasing the number of objects in the scene, in this experiment we are able to test not only whether participants are still able to predict in visually complex scenes, but also whether working memory plays a role in their ability to do so. As the average visual working memory capacity of humans is between 3 and 4 items (Luck & Vogel, 1997), we included a visual working memory task to determine whether participants with low working memory capacity show less pronounced anticipatory eye-movement behaviour.

### 3.1 Materials and Methods

#### 3.1.1 Participants

Twenty native speakers of Dutch (17 female, M_age_: 21.5 years, SD_age_: 1.76 years) were recruited from the Max Planck Institute for Psycholinguistics database. These participants had not participated in the previous experiment. The data of 27 participants was recorded, but six participants were discarded due to insufficient accuracy of the eye-tracking data and one stated during the debrief stage that they did not understand the virtual agent properly (clarity rating < 3 out of 5). The participants gave written informed consent prior to the experiment and were monetarily compensated for their participation.

#### 3.1.2 Materials and Design

The same materials and apparatus were used as described for Experiment 1. We selected four extra objects per scene that fit the theme of the scene (for example, a calculator in the office scene - see Appendix for a full list of added objects). The objects were not predictable given the restrictive verbs used in that scene, however, they were allowed to be candidates for completion in the unrestrictive conditions (i.e., “my colleagues hate it when someone throws away a - “). The objects were placed in realistic locations within each existing scene.

##### 3.1.2.1 Visual Working Memory Task

We used a saccadic sequential comparison task to assess visual working memory capacity. We chose this task as it arguably reflects the working memory used to complete the anticipatory language task in a reliable way: Participants view objects to be remembered and make a saccadic eye movement to the target object. The visual working memory task was performed after the participants completed the recall questionnaire and was conducted in the CAVE system - although the items were not rendered as 3D, the CAVE enabled us to use the eye-tracking system to record their eye movements and fixations. The visual working memory task took place on the middle screen only, with the entire array visible without the participant needing to move their head.

Our task is based on the one described by Heyselaar and colleagues (2011): Stimulus arrays consisted of sets of two to five coloured squares presented around a central fixation spot (Figure 3). For each set size, the spatial configuration of the squares remained identical across trials. For set size two, squares were on the right and left sides of the fixation spot. For set size three to five, squares were arranged equidistantly from each other with one square located directly above the fixation spot.

**Figure 3.**
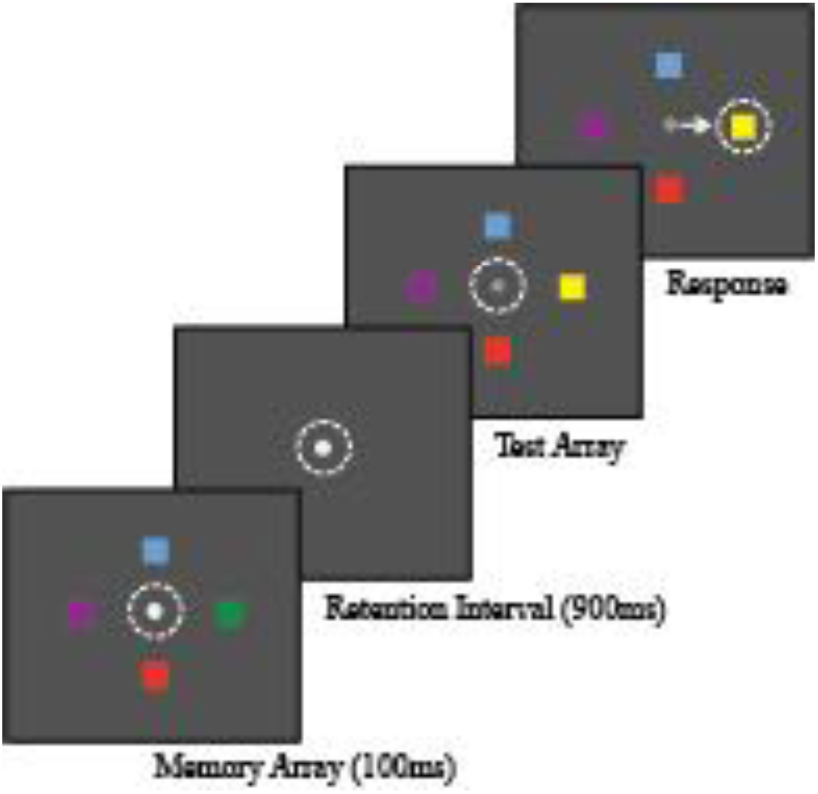
Depiction of a correctly performed trial in the sequential comparison task. Dotted lines and arrow represent current eye position and the saccade response. Participants were required to maintain fixation on the central fixation spot until the spot turned grey, signalling that they were allowed to move their eyes. The figure shows a correct trial. Adapted from Heyselaar et al., 2011.

The colour of each square was chosen randomly from a pre-determined library of six colors highly discriminable from each other. We used the Adobe Color Wheel (www.color.adobe.com) to choose six analogous colors. A given colour could only appear once in each array.

Figure 3 depicts the order of events in one trial. Each trial began with the presentation of a white fixation spot at the center of the middle screen. Participants were required to fixate this spot for a jittered period of 500 - 800ms. While they maintained fixation, a memory array composed of a randomly determined set of two to five squares was presented for 100ms. Offset of the memory array was followed by a 900ms retention interval, in which the display screen was blank with the exception of the central fixation spot. At the end of the retention interval, a test array was presented consisting of the same number and spatial configuration of the squares as in the memory array, but with the colour of one square changed. Concurrent with this, the fixation spot was dimmed and participants were required to make a saccade to the location of the changed square within 2 seconds. An inter-trial interval of a jittered 1000 - 1500ms followed before the next trial started.

Participants completed 80 trials, 20 for each set size. For each trial, there was always one square that was changed. The first square fixated was taken as the participant’s response. Therefore, participants could not fixate all squares within the 2s and still be marked as correct (unless the first square fixated was the changed square). This task took around 10 minutes to complete.

#### 3.1.3 Statistical analysis

For the main experiment, the same statistical analysis was used as described in Experiment 1. We removed 9.47% of all frames logged as object fixations and 4.70% of all frames logged as virtual agent fixations. To assess performance in the visual working memory task, we computed the proportion of correct responses.

### 3.2 Results

Participants were able to accurately identify which objects the virtual agent had named and which she had not 90.53% of the time (SD: 6.96%). Therefore we were confident that all participants listened to the virtual agent throughout the experiment.

Figure 4A illustrates the grand mean for this experiment. We again observed a robust increase in the proportion of looks to the target object after it was named. Figure 4B illustrates the proportion of looks per condition over time. Table 2 reports the summary output of the GAMM analysis.

**Figure 4.**
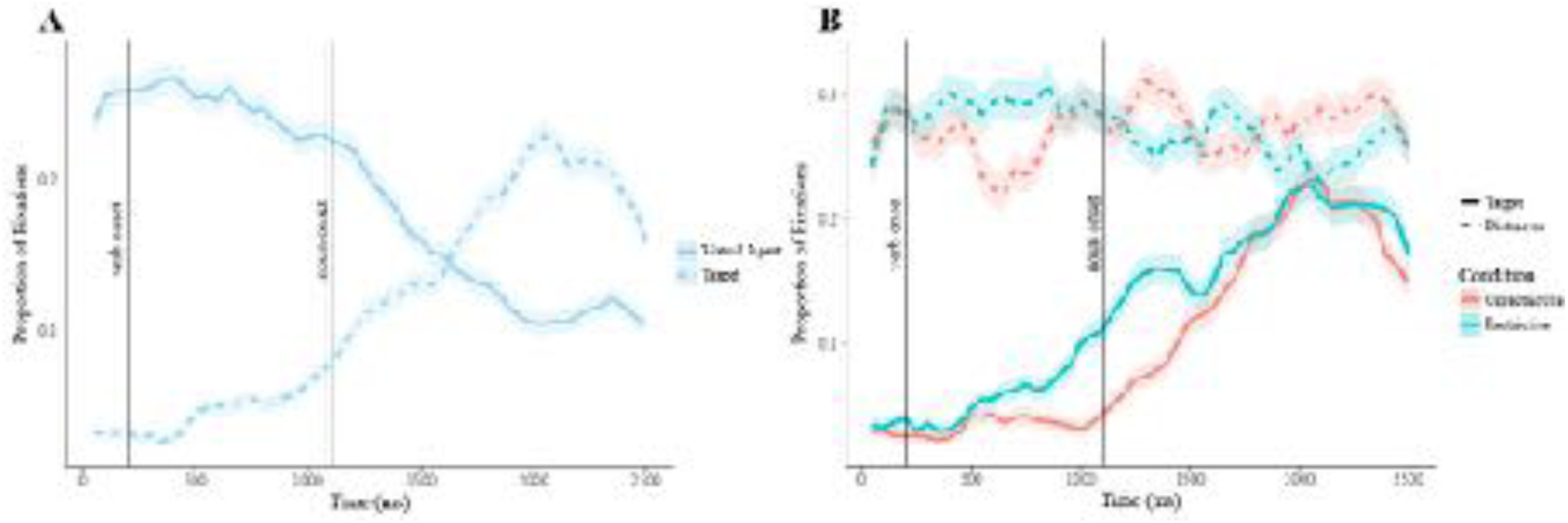
Mean proportions of fixations. **A**. To the target object and virtual agent. **B**. To the target and distractor objects shown per condition. Vertical lines indicate critical time points. The main statistical analysis was performed on the interval between verb onset and noun onset. Error clouds indicate standard error.

**Table 2.** Summary of the generalized additive mixed model for changes in target fixations over time, per condition (restrictive versus unrestrictive sentences). Effective degrees of freedom (edf), reference degrees of freedom (Ref.df)

| Parametric coefficients: |  |  |  |  |  |
| --- | --- | --- | --- | --- | --- |
|  | Estimate | SE | *t*-value | *p*-value |  |
| Intercept | -1.73 | 0.04 | -39.93 | < .001 | *** |
| Condition | -0.07 | 0.04 | -1.66 | .097 |  |
| Smooth terms |  |  |  |  |  |
|  | edf | Ref.df | *F*-value | *p*-value |  |
| Smooth for Time - Unrestrictive | 0.24 | 0.38 | 0.01 | .936 |  |
| Smooth for Time - Restrictive | 1 | 1 | 5.90 | .015 | * |
| Random effect for Subjects | 20.81 | 25.33 | 4.81 | < .001 | *** |
| Random effect for Sentences | 149.88 | 574 | 2.80 | < .001 | *** |
| *** < .001 |  |  |  |  |  |

We again observed an increase in the proportion of looks to the target object as a function of time for the restrictive (*p* = .015) but not the unrestrictive (*p* = .936) condition. The difference in the predicted values became significant between 688ms and 900ms after verb onset. No effect of condition on the proportion of distractor fixations was observed (*p* > .181).

We thus again observed anticipatory eye-movement behaviour for the restrictive condition versus the unrestrictive condition, in spite of an increase in the number of potential target objects. This suggests that even in scenes enriched with more objects, participants anticipated which object the virtual agent would name on the basis of restrictive information encountered at the verb.

#### 3.2.1 The Effect of Visual Working Memory Capacity on Anticipatory Behaviour

The median working memory capacity was 2.67 items (M: 2.663), in line with previous visual working memory capacity studies using sequential comparison (Luck & Vogel, 1997; Vogel & Machizawa, 2004; Vogel, Woodman, & Luck, 2006, inter alia). We therefore categorized participants as having low working memory (< 2.67) or high working memory (> 2.67). Figure 5 illustrates the fixation patterns of these two groups, per condition. Both groups showed anticipatory eye movements for the restrictive condition (*p* < .038), however, a model with difference smooths for the restrictive condition showed that participants with a higher working memory capacity fixate the target object earlier and more frequently compared to the low working memory capacity group (*p* = .009).

**Figure 5.**
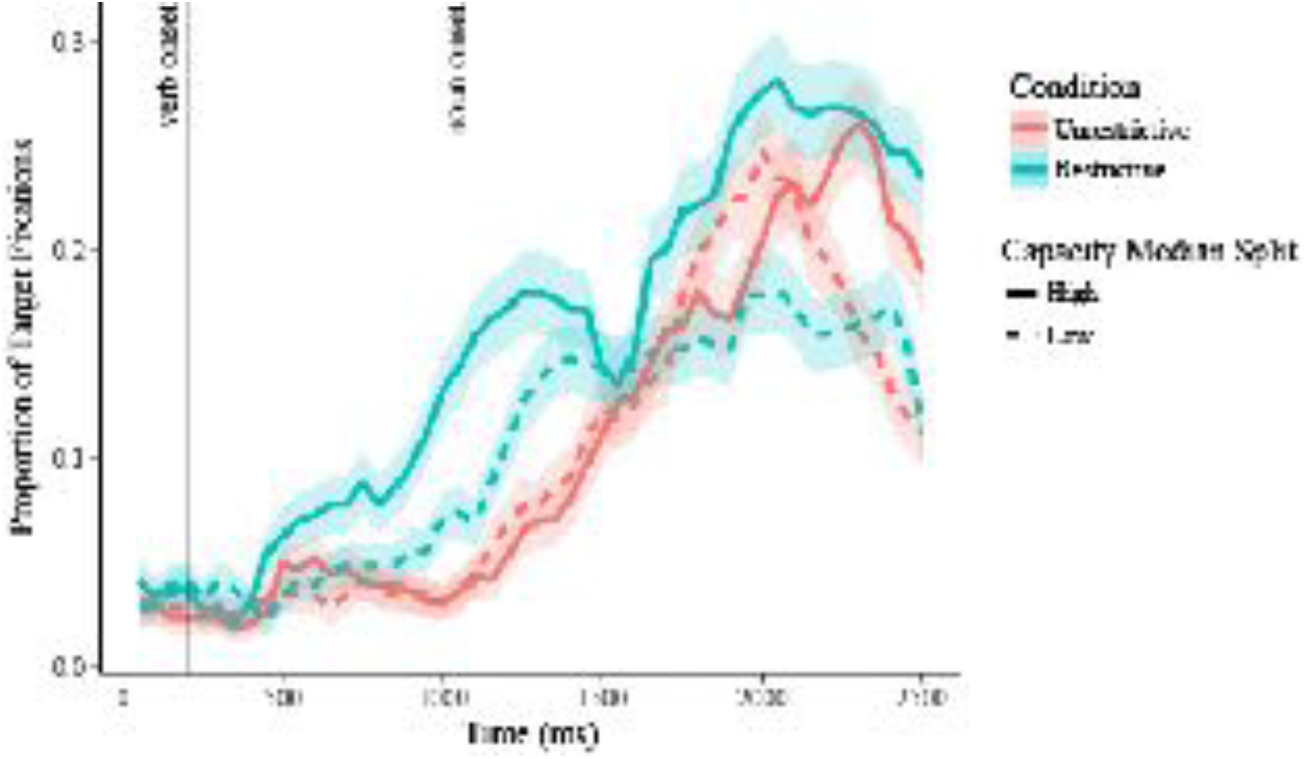
Mean proportions of target fixations for participants with low (< 2.67 items) and high working memory capacity (> 2.67 items), per condition. Participants with high working memory capacity showed earlier and more robust anticipatory eye-movements compared to the low working memory capacity participants *(p* = .009), although both groups showed significant anticipatory behaviour relative to the unrestrictive condition *(p* < .038). Error clouds represent standard error.

## 4. Experiment 3: Peripheral objects

The results from Experiment 2 suggest that working memory does indeed play a role in anticipatory eye-movement behaviour, as proposed by Huettig and colleagues (2011). They suggest that participants maintain representations of the objects in the scene in working memory, which allows for a quick selection once the restrictive verb is processed. If this is indeed how working memory is involved in anticipatory eye-movement behaviour (and perhaps linguistic prediction in general), then if participants have not encoded the object into working memory, the object should not be considered a potential target when the restrictive verb is processed.

We tested this theory by placing the target objects in the visual periphery of the participants (the left and right screen as illustrated in Figure 1, with two target objects on each side). This essentially hides the objects from the participant, although they are clearly identifiable if the participant turns their head. If participants indeed only consider objects already maintained in working memory, then only objects viewed during the preview time will be fixated. However, if another mechanism is responsible, such as participants initiating a visual search for a potential object upon hearing a potentially restrictive verb, then we should still see increased fixations on the target object, although perhaps later in time.

### 4.1 Materials and Methods

#### 4.1.1 Participants

Twenty native speakers of Dutch (17 female, M_age_: 22.4 years, SD_age_: 2.44 years) were recruited from the Max Planck Institute for Psycholinguistics database. These participants had not participated in the previous experiments. The data of 24 participants was recorded, but three participants were discarded due to insufficient accuracy of the eye-tracking data and one stated during the debrief stage that they did not understand the virtual agent properly (clarity rating < 3 out of 5). The participants gave written informed consent prior to the experiment and were monetarily compensated for their participation.

#### 4.1.2 Materials

The same materials were used as described for Experiment 1. Only the location of the four target objects per scene was changed such that two were present on each of the peripheral screens.

#### 4.1.3 Statistical analysis

The same statistical analysis was used as that described in Experiment 1. We removed 11.85% of all frames logged as object fixations and 5.18% of all frames logged as virtual agent fixations.

### 4.2 Results

Participants were able to accurately identify which objects the virtual agent had named and which she had not named 93.16% of the time (SD: 4.78%). Therefore we are confident that all participants listened to the virtual agent throughout the experiment.

Figure 6A illustrates the grand mean for this experiment. We did not see the robust increase in the proportion of looks to the target object that we observed in the other experiments. In fact, the peak (0.13) occurred 976ms after sentence offset. This suggests that some participants did search for the object, even after it was named; however, the majority did not. Only 37.42% of the target objects had been fixated by the participants before they were named by the virtual agent. However, even for these fixated objects (229 trials), we still see no anticipatory eye movements. Perhaps as the objects were not in the shared space between the participant and the avatar (whereas they were in Experiment 1), they were encoded differently (or perhaps not all) and hence not considered as a potential target in the upcoming sentence.

**Figure 6.**
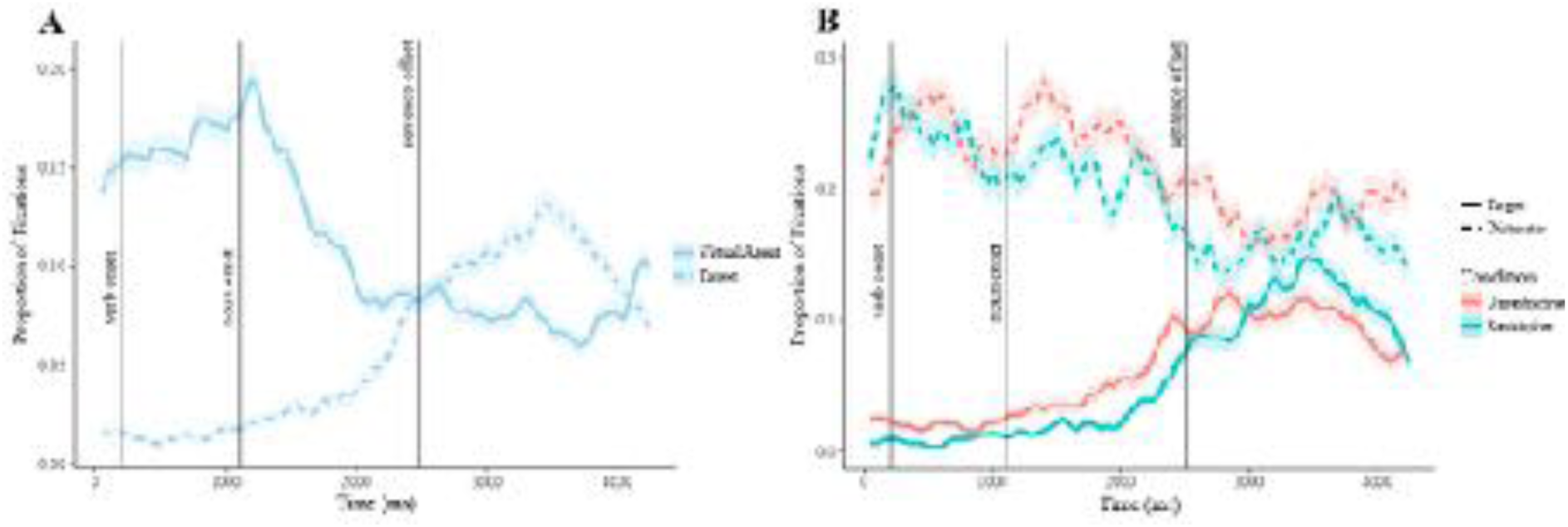
Mean proportions of fixations. **A**. To the target object and virtual agent. **B**. To the target and distractor objects shown per condition. Vertical lines indicate critical time points. The main statistical analysis was performed on the interval between verb onset and noun onset. Error clouds indicate standard error.

Figure 6B illustrates the proportion of looks per condition. Table 3 reports the summary output of the GAMM analysis.

**Table 3.**
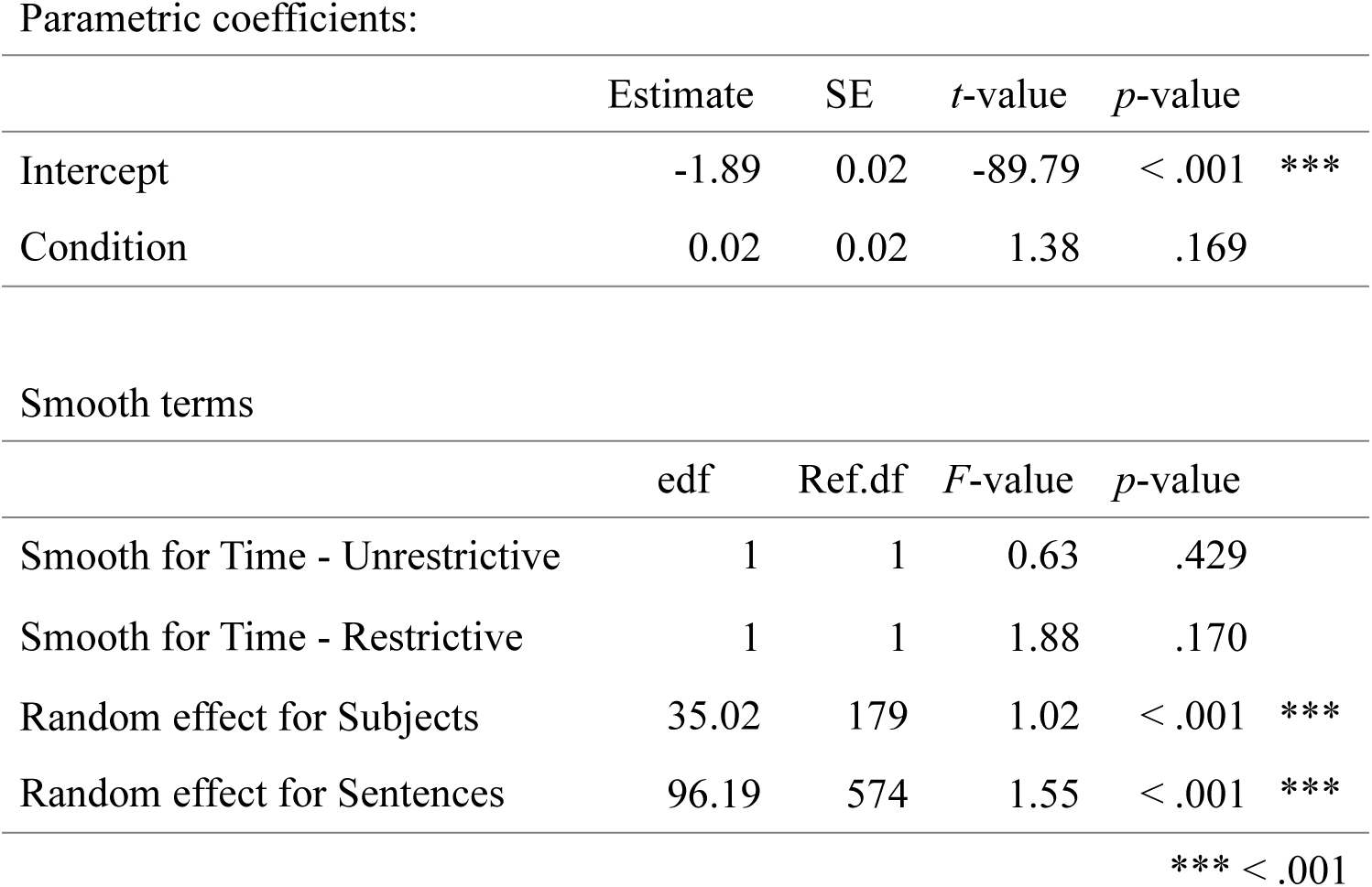
Summary of the generalized additive mixed model for changes in target fixations over time, per condition (restrictive versus unrestrictive sentences). Effective degrees of freedom (edf), reference degrees of freedom (Ref.df)

As illustrated in Figure 6B, there were no anticipatory looks to the target object during the critical window. One might argue that, due to the target objects being present in the periphery, the 900ms window was not enough time to fixate the object before it was named. This would mean that past the critical window, the peak target fixations should be earlier for the restrictive compared to the unrestrictive condition. However, it is clear from the figure that this was also not the case.

Therefore, if the target object is not in the immediate view of the participant, no anticipatory eye movements are made towards it if the verb is restrictive. This suggests that 1) hearing a restrictive verb does not initiate a visual search from the participant to look for an object that could fit that verb, and therefore 2) anticipatory behaviour in the visual world paradigm only occurs when an object has been viewed before the respective restrictive sentence is spoken. In line with our working memory capacity findings from Experiment 2, this indicates that objects are held online once viewed, allowing for a quick selection once a restrictive verb is comprehended. If a verb is heard and no viable object is online, no visual search is initiated in an attempt to find a candidate before or after the noun is encountered.

## 5. Experiment 4: Manipulating referent predictability

Experiments 1 and 2 have shown that participants show anticipatory eye movement behaviour during restrictive sentences even when faced with rich everyday scenes including 10 potential referent objects. This could be because every sentence spoken by the virtual agent concerned an object in the scene, a pattern that participants could have realized early in the experiment. Therefore, in Experiment 4 we introduced eight filler sentences per scene: Sentences that did not concern objects present in the scene. These filler sentences were similar to the restrictive/unrestrictive sentences in that they did concern an object (e.g., “People bring their own briefcase to work”) and therefore participants were not able to detect whether a sentence spoken by the virtual agent was a filler or not until the object was named. Verbs were again controlled to ensure that they were not predictive of objects already present in the scene.

In sum, in this experiment only 50% of all sentences spoken concerned an object that the participants could fixate, and in only 25% of all sentences spoken, the target object could be anticipated given the verb.

### 5.1 Materials and Methods

#### 5.1.1 Participants

Twenty native speakers of Dutch (12 female, M_age_: 22.7 years, SD_age_: 2.11 years) were recruited from the Max Planck Institute for Psycholinguistics database. These participants had not participated in the previous experiments. Data from one additional participant was discarded due to insufficient accuracy of the eye-tracking data. The participants gave written informed consent prior to the experiment and were monetarily compensated for their participation.

#### 5.1.2 Materials

The same materials were used as described for Experiment 1. We created eight extra filler sentences per scene (64 extra sentences in total). Frequency of the verbs between the three conditions (restrictive, unrestrictive, and filler) was not significantly different (*F*(2,127) = 1.861, *p* = .160) although length was (*F*(2,127) = 8.12, *p* < .001). *Post-hoc* comparison showed that the filler verbs were significantly longer (M = 7.59 characters, SD = 2.32, Tukey’s HSD, *p* < .033) compared to the restrictive (M = 5.91 characters, SD = 1.69) and unrestrictive (M = 6.47 characters, SD = 1.78) conditions.

Sentences from all three conditions were presented randomly in each scene. Due to the increase in sentences, the task took 15 minutes to complete.

#### 5.1.3 Statistical analysis

The same statistical analysis was used as that described in Experiment 1. We removed 7.67% of all frames logged as object fixations and 2.32% of all frames logged as virtual agent fixations.

### 5.2 Results

Participants were able to accurately identify which objects the virtual agent had named and which she had not 82.68% of the time (SD: 8.67%). Therefore we are confident that all participants listened to the virtual agent throughout the experiment.

Figure 7A illustrates the grand mean for this experiment. We again observed a robust increase in the proportion of looks to the target object after it is named. Figure 7B illustrates the proportion of looks per condition. Table 4 reports the summary output of the GAMM analysis. For this analysis, the filler condition was not included as, by definition, there was no target object to fixate.

**Figure 7.**
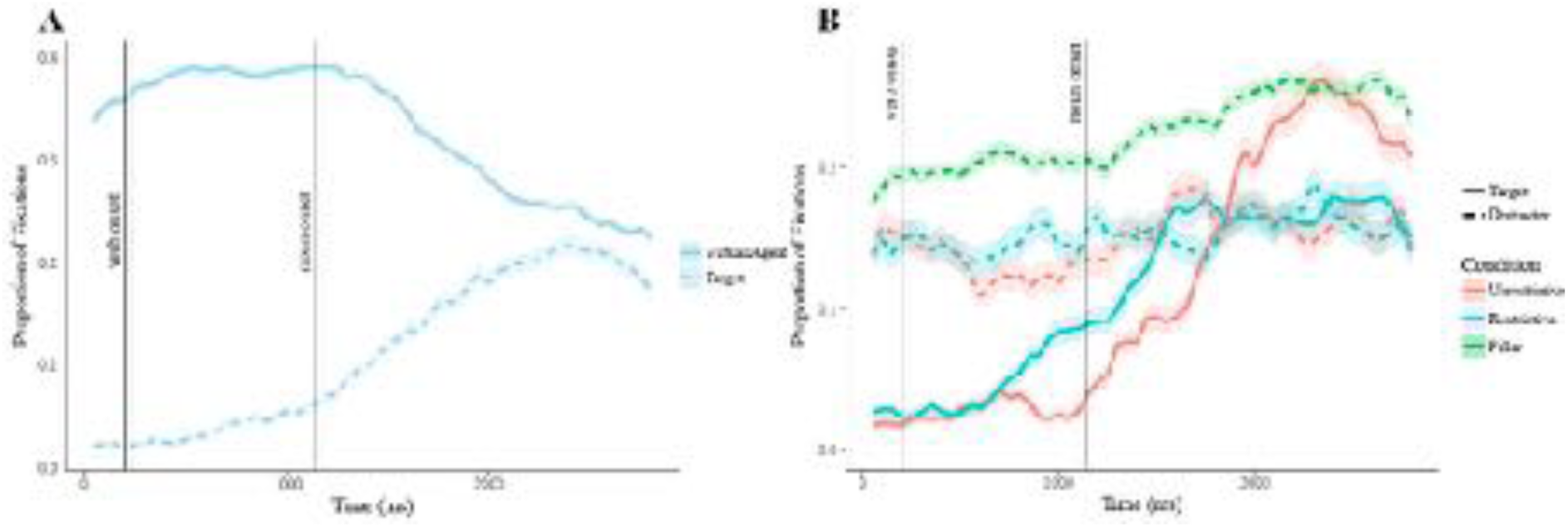
Mean proportions of fixations. **A**. To the target object and virtual agent. **B**. To the target and distractor objects shown per condition. Vertical lines indicate critical time points. The main statistical analysis was performed on the interval between verb onset and noun onset. Error clouds indicate standard error.

**Table 4.** Summary of the generalized additive mixed model for changes in target fixations over time, per condition (restrictive versus unrestrictive sentences). Effective degrees of freedom (edf), reference degrees of freedom (Ref.df)

| Parametric coefficients: |  |  |  |  |  |
| --- | --- | --- | --- | --- | --- |
|  | Estimate | SE | *t*-value | *p*-value |  |
| Intercept | -1.77 | 0.04 | -46.17 | < .001 | *** |
| Condition | -0.05 | 0.03 | -1.50 | .133 |  |
| Smooth terms |  |  |  |  |  |
|  | edf | Ref.df | *F*-value | *p*-value |  |
| Smooth for Time - Unrestrictive | 1.96 | 2.33 | 1.00 | .458 |  |
| Smooth for Time - Restrictive | 1 | 1 | 19.84 | < .001 | *** |
| Random effect for Subjects | 41.25 | 179 | 1.30 | < .001 | *** |
| Random effect for Sentences | 157.12 | 574 | 2.40 | < .001 | *** |

\*\*\* < .001

We again observed an increase in the proportion of looks to the target object as a function of time for the restrictive (*p* < .001) but not the unrestrictive (*p* = .458) condition. The difference in the predicted values became significant between 710ms and 900ms after verb onset. We observed no effect of condition on the proportion of distractor fixations for any of the three conditions (restrictive: *p* = .551; unrestrictive: *p* = .646; filler: *p* = .716).

Thus, we again observed significant anticipatory eye-movement behaviour for the restrictive condition, even though this behaviour was only efficient for 25% of the sentences heard.

## 6. Overall Results

For a comparison of the four experiments, Figure 8 illustrates the looks to the target object in the restrictive condition only, for the four experiments. As there was no significant anticipatory eye-movement behaviour for Experiment 3, we conducted a GAMM analysis of the remaining three experiments (Experiments 1, 2, and 4) to determine whether the observed anticipatory eye-movements in Experiments 2 and 4 were significantly different from those observed in Experiment 1. The model confirmed that participants showed more robust anticipatory eye-movement behaviour for Experiment 1 compared to Experiments 2 and 4 (*p =* .006).

**Figure 8.**
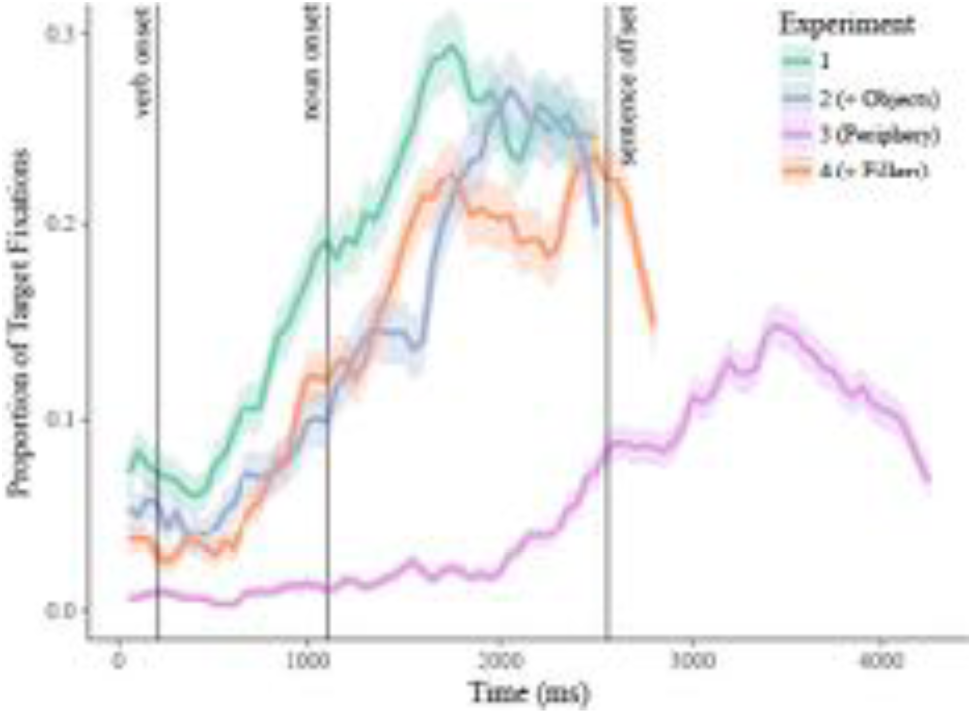
Mean proportions of fixations per experiment to the target object during the restrictive condition. Vertical lines indicate critical time points. Experiment 1 induced significantly more fixations to the target object for both plots *(p* < .007). Error clouds represent standard error.

## 7. Discussion

Prediction is commonly considered a central component of cognition. When processing incoming language input, the fact that we may predict upcoming words is generally used to explain why conversation and turn-taking are such efficient linguistic feats. In four virtual reality experiments, we here tested whether a well-established marker of linguistic prediction (i.e. anticipatory eye movements as observed in the visual world paradigm) replicated when increasing the naturalness of the paradigm by means of i) immersing participants in naturalistic everyday scenes, ii) increasing the number of distractor objects present, iii) manipulating the location of referents in central versus peripheral vision, and iv) modifying the proportion of predictable noun-referents in the experiment. After all, previous experimental studies have mainly shown that listeners *can* predict, not necessarily that they *do* predict in more naturalistic everyday settings.

In the current study we thus used anticipatory eye-movements as a measure of prediction (Altmann & Kamide, 1999). If participants predict the upcoming referent in naturalistic situations, we would expect robust anticipatory eye-movements towards the referent object after participants heard the restrictive verb (i.e. when the target object was identifiable based on the verb alone) compared to the unrestrictive verb (when the target object could not be identified based on verb information alone). Thus if participants fixate the referent object significantly more and earlier after the verb was spoken but before the object was named in the restrictive condition, we would interpret that as evidence for predictive processing. This is exactly what we found in three of our four experiments. We were thus able to replicate the behaviour seen in traditional 2D (e.g., Altmann & Kamide, 1999) and 3D (e.g., Eichert et al., 2017) versions of the visual world paradigm. We will now describe in detail the findings and contributions of each of our four experiments.

### 7.1 Prediction in Naturalistic Environments

The main aim of the current study was to determine whether we still predict in naturalistic everyday scenes by increasing the ecological validity of the visual world paradigm (VWP). In Experiment 1, we diverged from the traditional visual world paradigm methodology by increasing the number of objects per scene (6 instead of 4), increasing the number of sentences per scene (4 instead of 1), and having a virtual agent deliver these sentences to the participants. As stated above, we were able to replicate anticipatory eye-movements in these situations, and thus for the remainder of the studies we continued to increase the number of objects (Experiment 2) and sentences (Experiment 4) to test whether participants still anticipated upcoming language input in these situations.

In Experiment 2, we increased the number of objects present to 10. Previous studies have presented conflicting evidence on whether increased visual complexity should influence the anticipatory behaviour of participants (Andersson et al., 2011; Coco & Keller, 2015 versus Sorensen & Bailey, 2006), and hence it was up for debate whether we would observe language-driven eye-movements in everyday settings. In our Experiment 2, however, participants still showed robust anticipatory eye-movement behaviour, albeit the time window of significance started 374ms later compared to Experiment 1.

In Experiment 4 we increased the realism of the VWP by manipulating the probability of hearing a predictable sentence. This was done in two ways: 1) 50% of the sentences did not refer to objects in the scene (filler sentences) which ensured that 2) only 25% of sentences heard included a restrictive verb. Even under these conditions, in which predictions of specific upcoming noun referents were confirmed in only 25% of the cases, participants still exhibited robust and significant anticipatory eye-movements towards the referent objects in the restrictive compared to the unrestrictive conditions.

The results from Experiment 2 and 4 have provided evidence that participants anticipate upcoming linguistic information in visually complex scenes, and even when it is only efficient to do so 25% of the time. Additionally, our experiments allowed us to test two mechanisms that play important roles in current theories of prediction but to date have little to no empirical evidence to support their contributions: Working memory and error-based learning.

### 7.2 The Role of Working Memory

In Experiment 3, we placed the target objects in the participants’ peripheral field of view, to determine whether hearing a restrictive verb would initiate a visual search for that object. Even after the conclusion of the sentence, the majority of the target objects (62.5%) had not been fixated, suggesting that participants did not search for the object even after it was named by the virtual agent. However, for the 37.42% of the objects that were viewed before the sentence began, no anticipatory eye movements were seen either. As the placement of the objects in the periphery was the only difference in this experiment compared to Experiment 1, the new locations of the objects seems to have changed the way participants encode and/or remember them. This suggests that anticipatory eye-movements are only executed for objects the participants have i) already viewed and ii) encoded as relevant to the current context. This is in line with studies suggesting that anticipatory eye movements depend on mental representations the listeners construct about the environment (Altmann & Kamide, 2009; Knoeferle & Crocker, 2006, 2007; cf. Huettig et al., 2011). An object the participants have not seen and/or has not been tagged as potentially relevant would indeed not necessarily form part of their mental representation.

Huettig and colleagues (2011) argue that anticipatory eye movements in the VWP are supported by working memory: Objects in the display are first encoded in a visuospatial type of working memory (cf. Baddeley, 1998; Cavanagh & Alvarez, 2005; Pylyshyn, 1989), which triggers perceptual hypotheses in long-term memory. These hypotheses then trigger a cascade of activations of associated semantic and phonological codes, all within a few hundreds of milliseconds (cf. Huettig & McQueen, 2007). This results in a nexus of associated knowledge, which is bound to an object’s location within working memory. However, working memory has a limited capacity: The capacity of visual working memory has been shown to be between 3 and 4 unrelated items (coloured squares on a black background; Luck & Vogel, 1997; Vogel & Machizawa, 2004). In VWP tasks, the items are usually related to and embedded in an environment, in which items are meaningfully connected to the total scene. Hence it is possible that more than 4 items can be retained in working memory.

By increasing the number of objects to 10 (as we did in Experiment 2), we taxed the visual working memory system and hence should measure a decrease in anticipatory eye-movements. This is indeed what we observed: Although we still found a robust increase in anticipatory eye-movements for the restrictive versus unrestrictive condition, it was significantly lower compared to Experiment 1. When splitting the participants by working memory capacity, we observed that participants with a higher working memory capacity fixated the target object earlier and more frequently compared to the low working memory capacity group. These results are in support of a role of working memory in prediction.

The proposal for a role of working memory in anticipating linguistic information is not new (Huettig, Olivers, et al., 2011; Knoeferle & Crocker, 2007), although there have only been two studies (to our knowledge) that have attempted to provide empirical evidence to support this proposal. Huettig and Janse (2016) found a positive correlation between the ratio of target-distractor object looks and a working memory construct score, such that participants with a higher working memory construct score showed a stronger prediction tendency. A recent study by Ito and colleagues (Ito, Corley, & Pickering, 2018) provided more causal evidence by demonstrating that participants showed reduced anticipatory looks to the referent object if they were required to simultaneously remember five words.

We build upon these two studies by being the first to provide a direct link between a participant’s working memory capacity and their anticipatory eye-movement behaviour.

### 7.3 Error-based Learning

In Experiment 4 we manipulated referent predictability by having only 25% of the sentences contain a verb that could be used to predict the specific upcoming referent. The remaining sentences were either unrestrictive (25%) or did not refer to an object present in the scene (50%). Many theories propose that prediction is supported by an error-based learning mechanism: We anticipate upcoming linguistic information based on past experiences (Chang, 2002; Dell & Chang, 2014). If this were the case, then one would expect that participants stop exhibiting anticipatory eye-movements as this would be inefficient given the statistical probability of an object being either present in the scene or predictable given the verb. Surprisingly, this is not what we observed: Participants still fixated the target object earlier and more frequently in the restrictive compared to the unrestrictive condition. However, when compared to Experiment 1, where all sentences refer to an object in the scene and 50% of the sentences heard include a restrictive verb, the magnitude of the anticipatory effect was significantly smaller (*p* < .006).

In our experiments we only measured looks to the target object, and thus there may have been other changes in behaviour that we missed. For example, Hintz and Huettig (2015) showed a change in fixation behaviour in more complex scenes. In the simpler experiments (simple line drawings in a grid), they showed that participants fixated on semantic and phonological competitors as the sentence unfolds. However, when conducting the experiment with more complex scenes (introducing characters that interact with the objects), they did not see any shifts towards phonological competitors, supporting the claim that the word-object mapping preferences were different between the two scene types.

### 7.4 Conclusion

Do we predict upcoming linguistic content in rich, naturalistic environments? The evidence provided here is mixed. On the one hand, we observed robust anticipatory eye movements in naturalistic scenes, even when these scenes contained a relatively large number of objects and a relatively small number of sentences that allowed for a predicted noun-referent to be confirmed by the speaker’s unfolding speech. These findings are in line with the proposed role of working memory in supporting our cognitive prediction machinery. On the other hand, however, the well-established effect of prediction disappeared when referent objects were placed in participants’ peripheral vision. This restricts the generalizability of the current and earlier prediction findings to situations in which interlocutors have actively encoded the objects in their immediate environment before communicating about them.

## Appendix

The verbs (underlined) are written restrictive/unrestrictive. The target object is boldfaced. Italicized sentences are the translated versions of the Dutch stimuli.

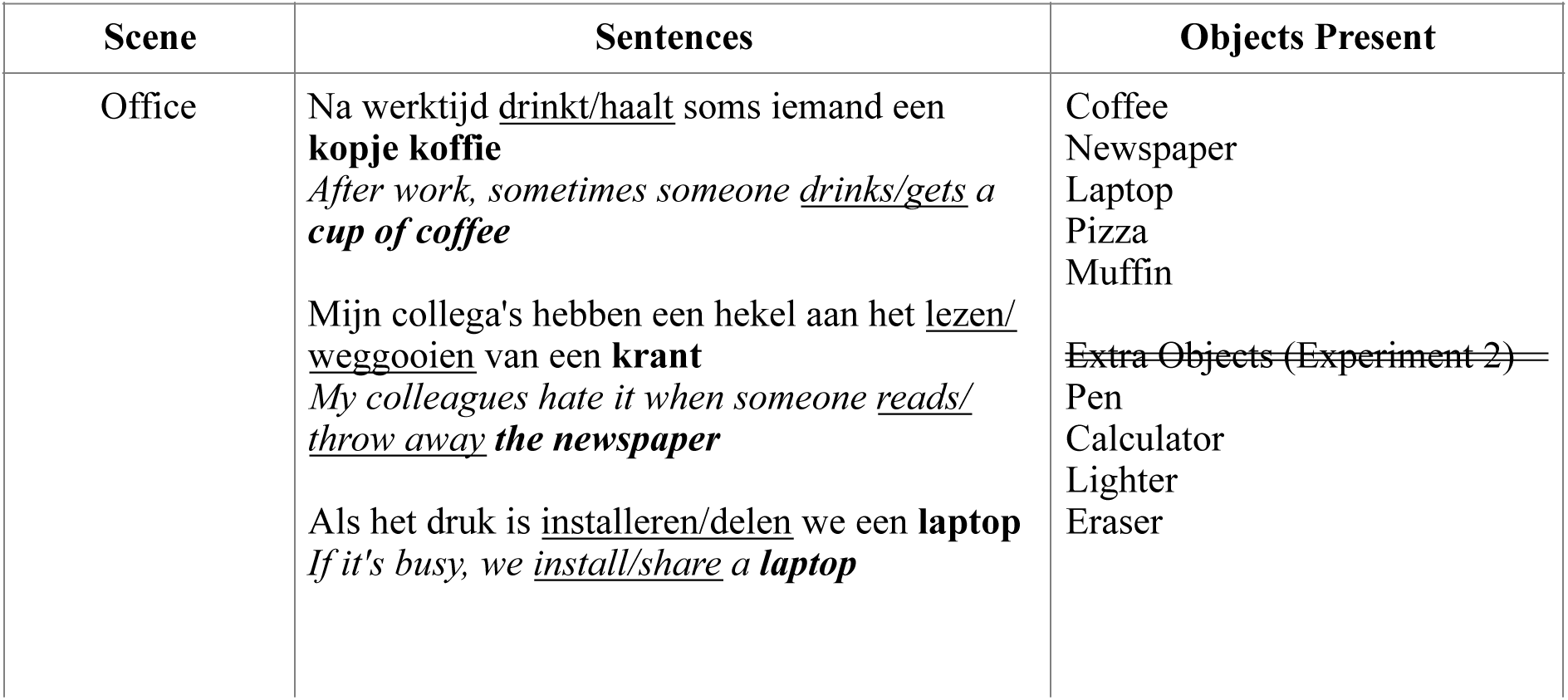
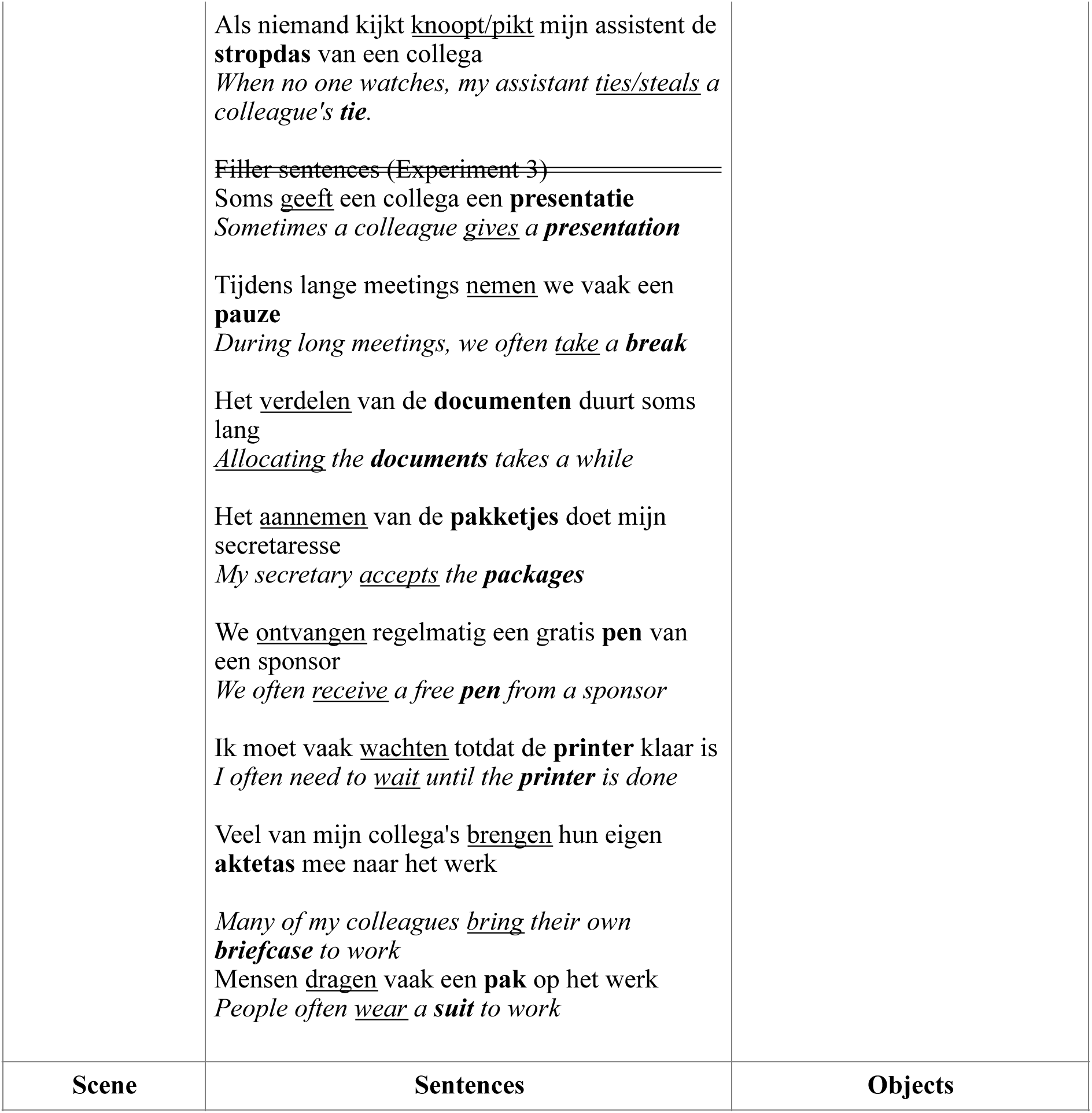
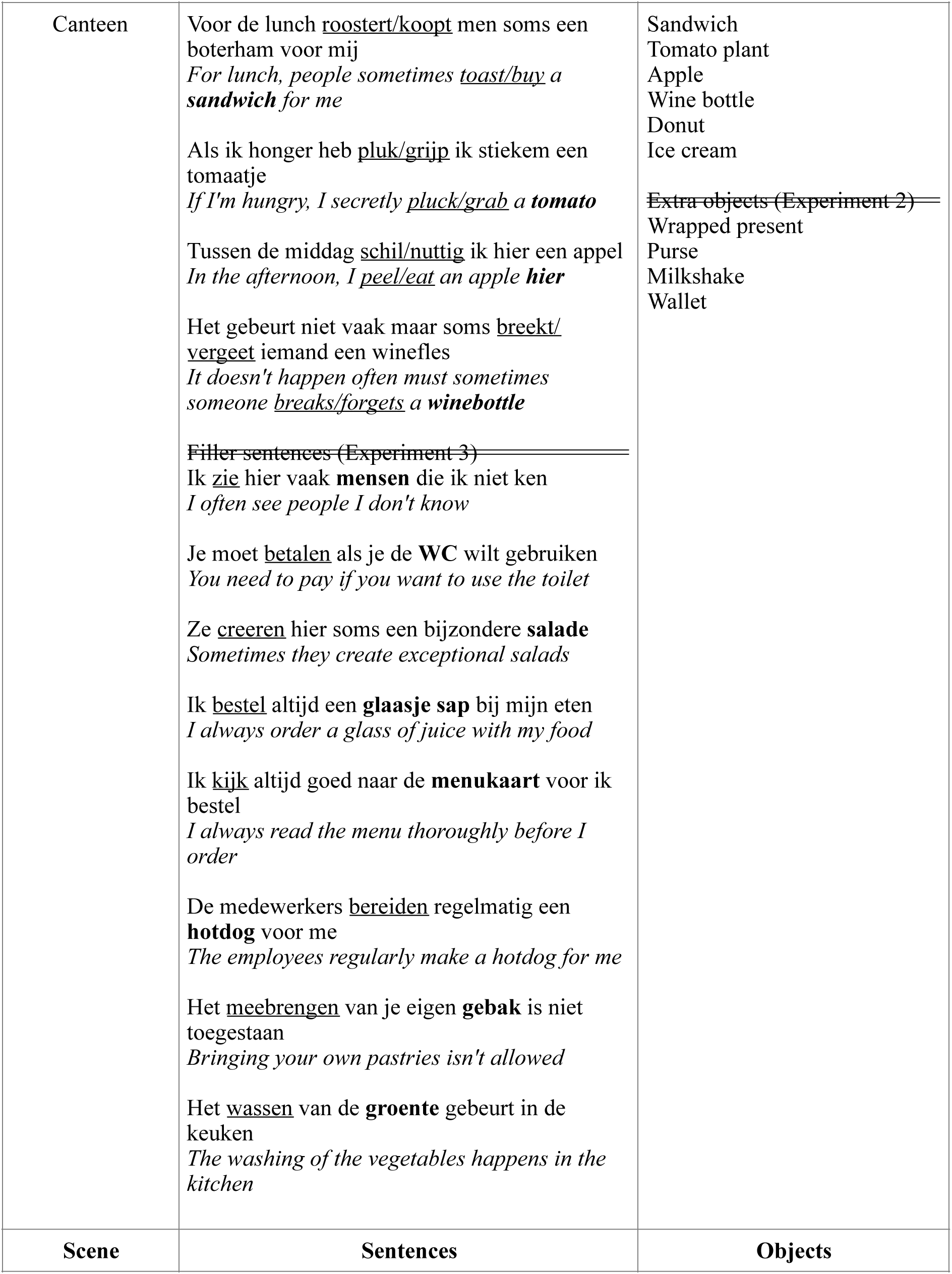
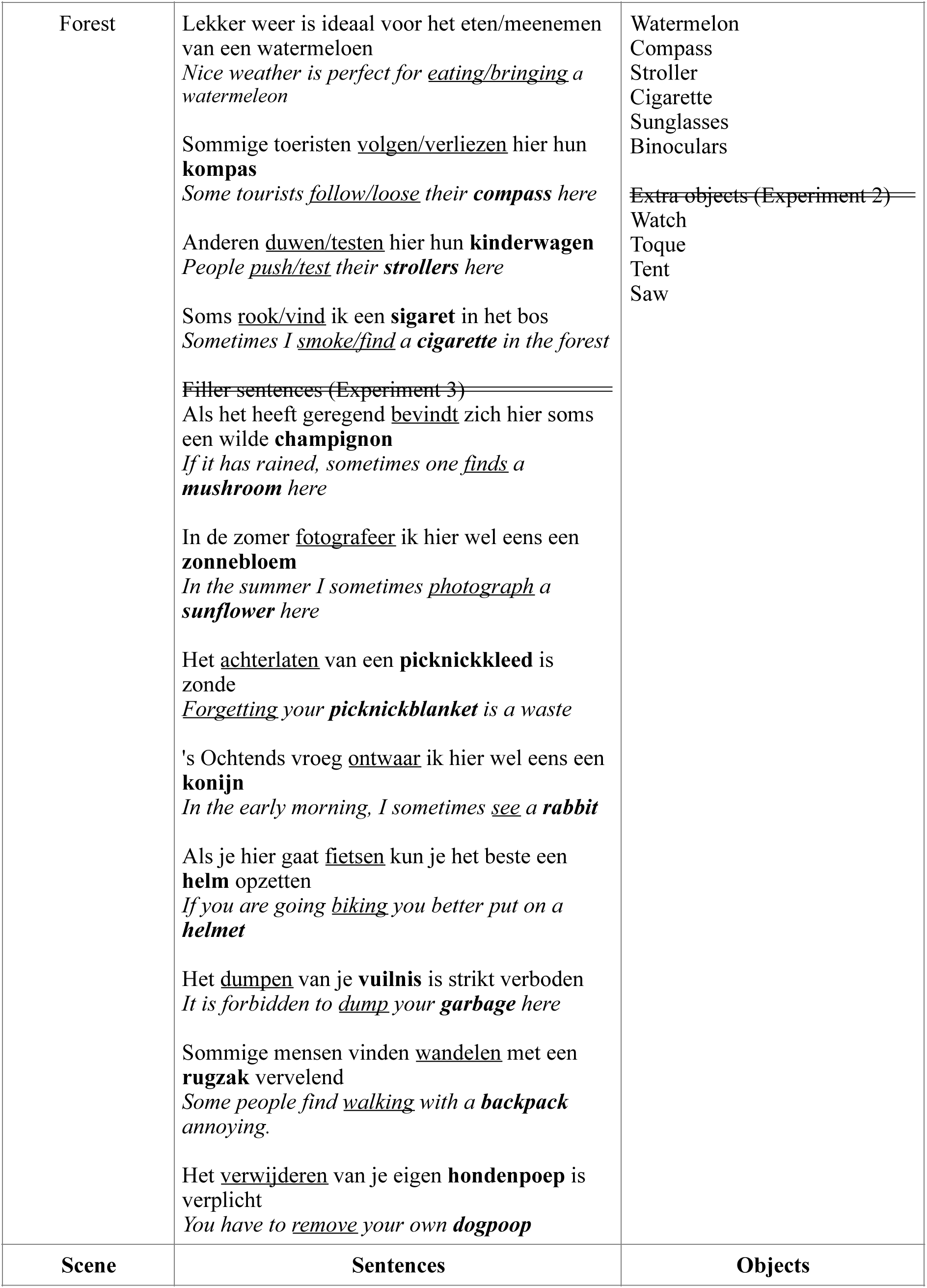
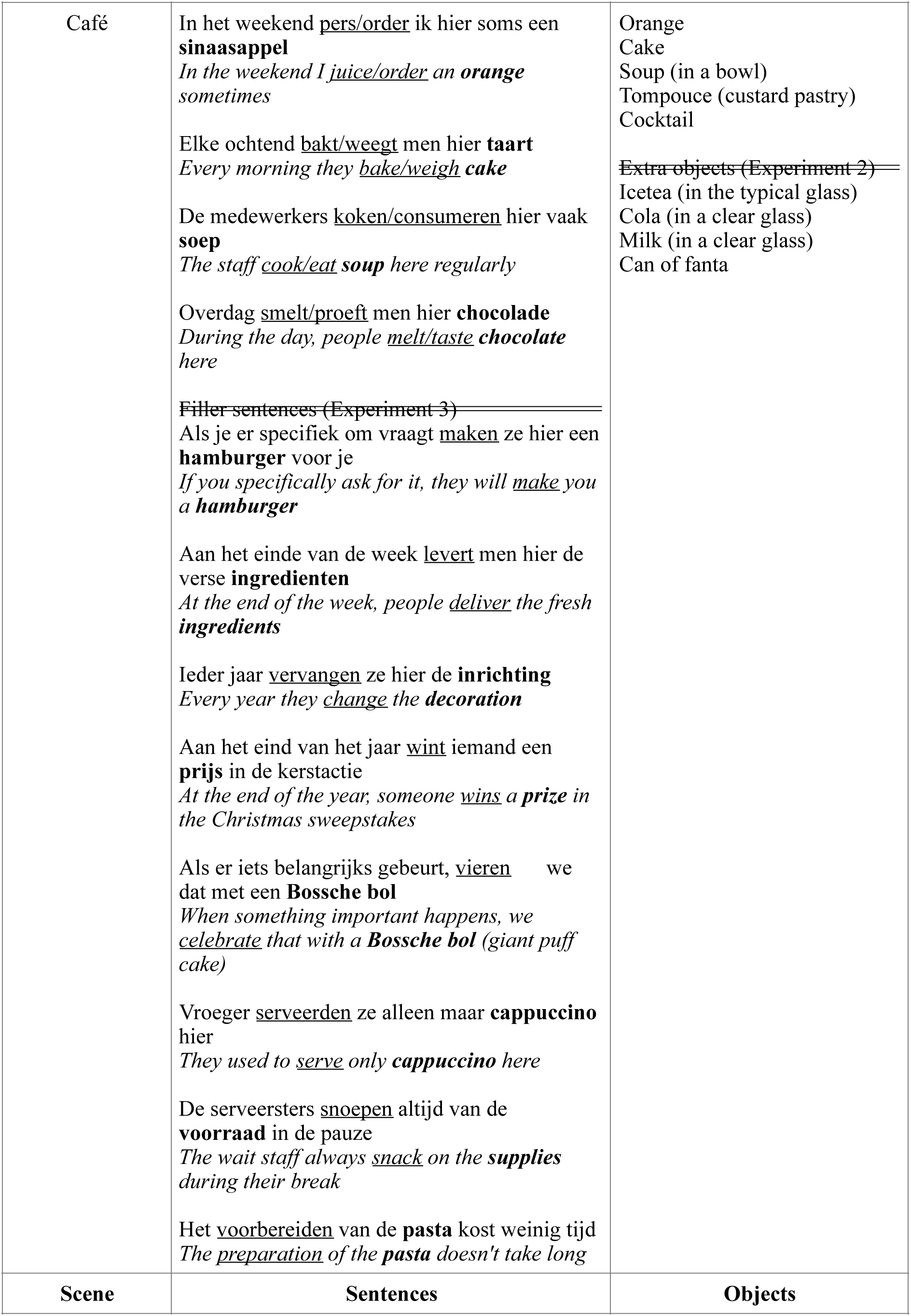
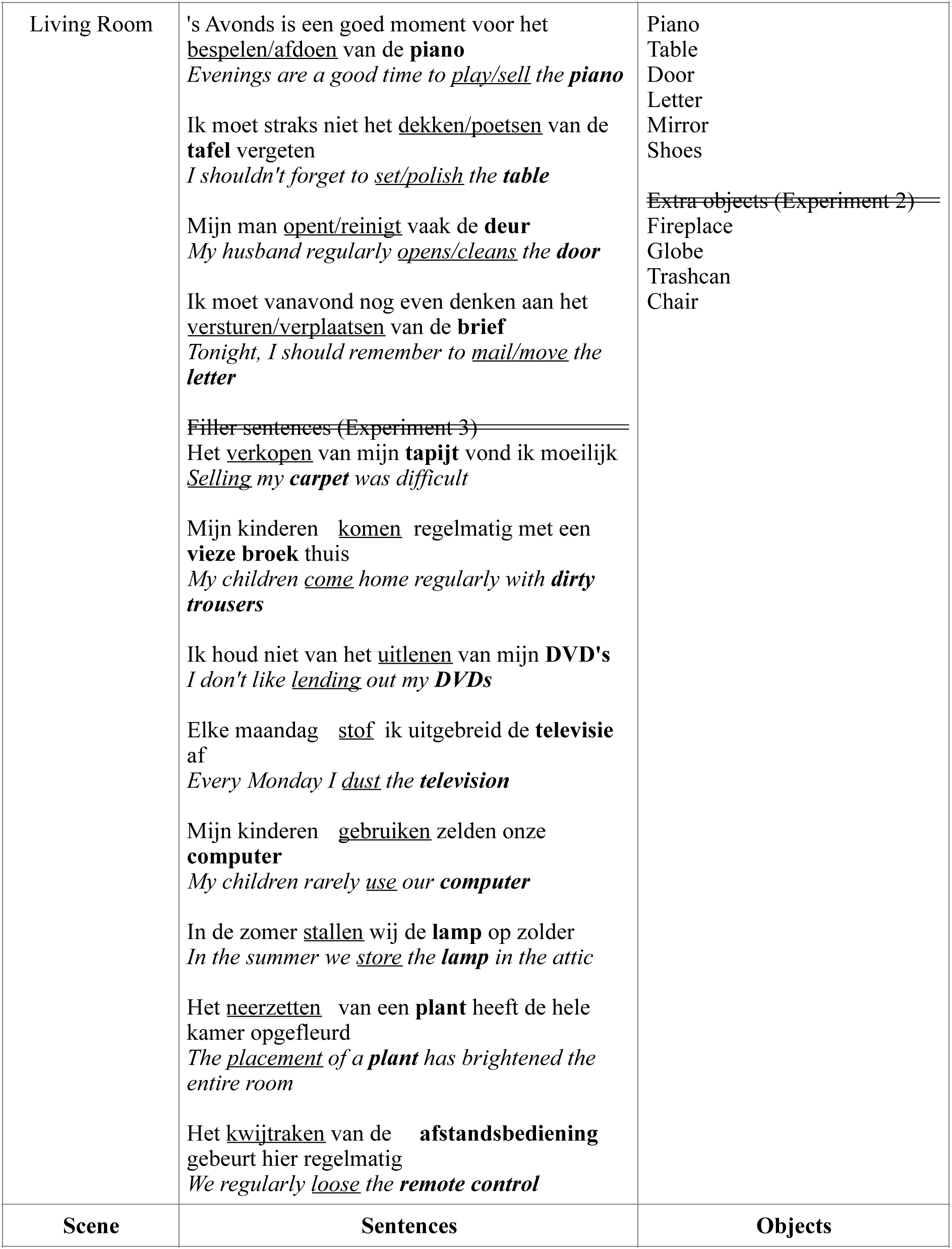
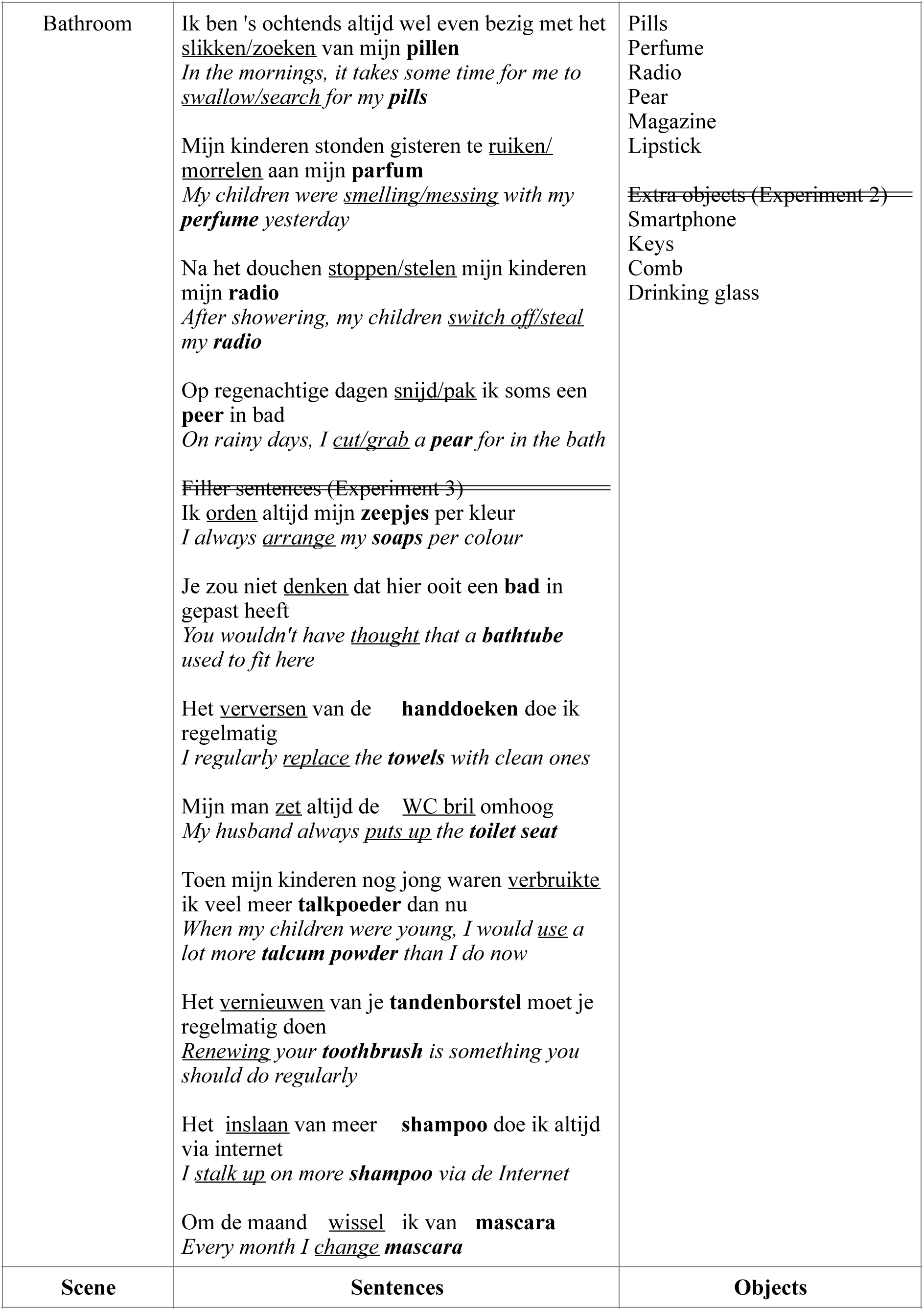
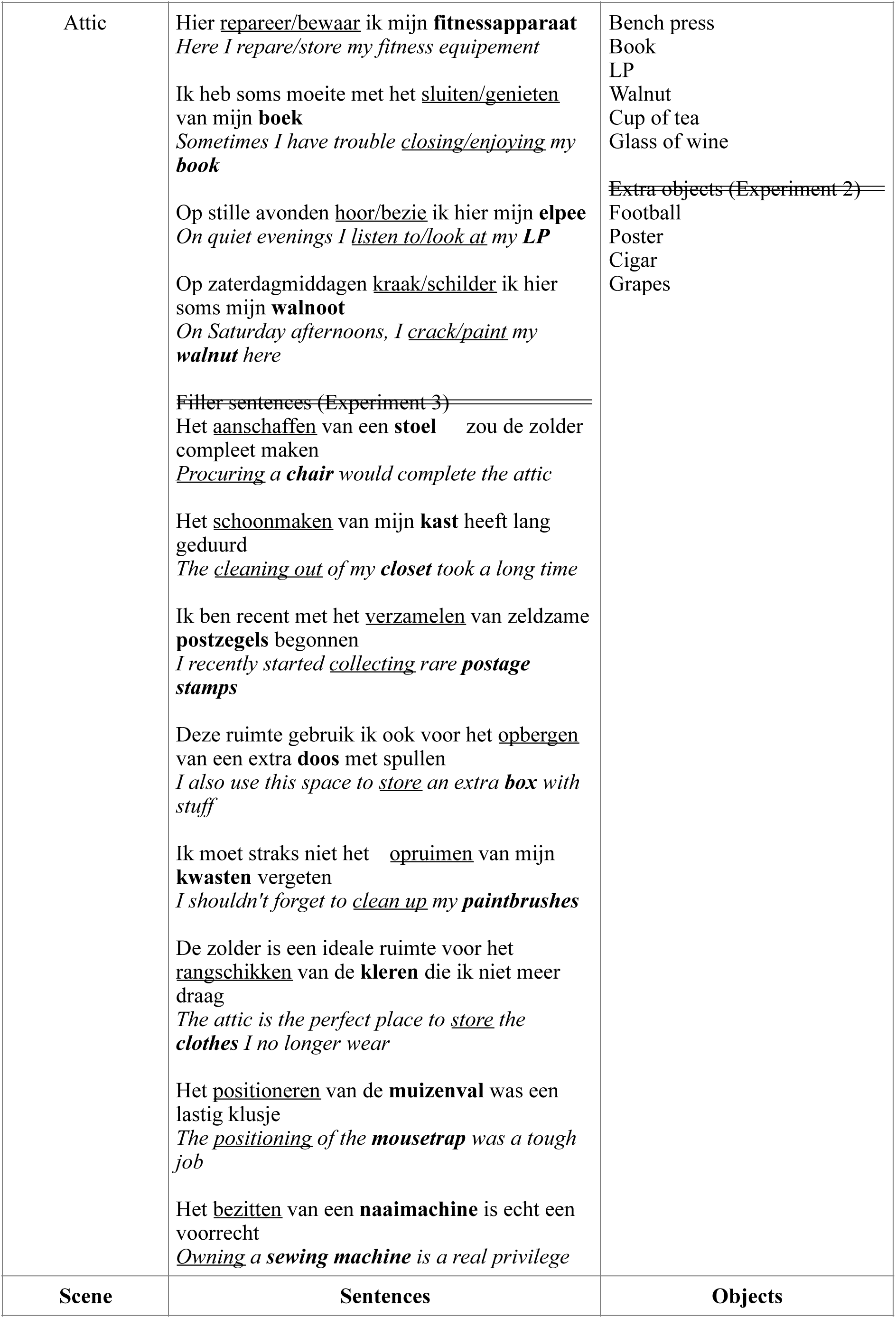
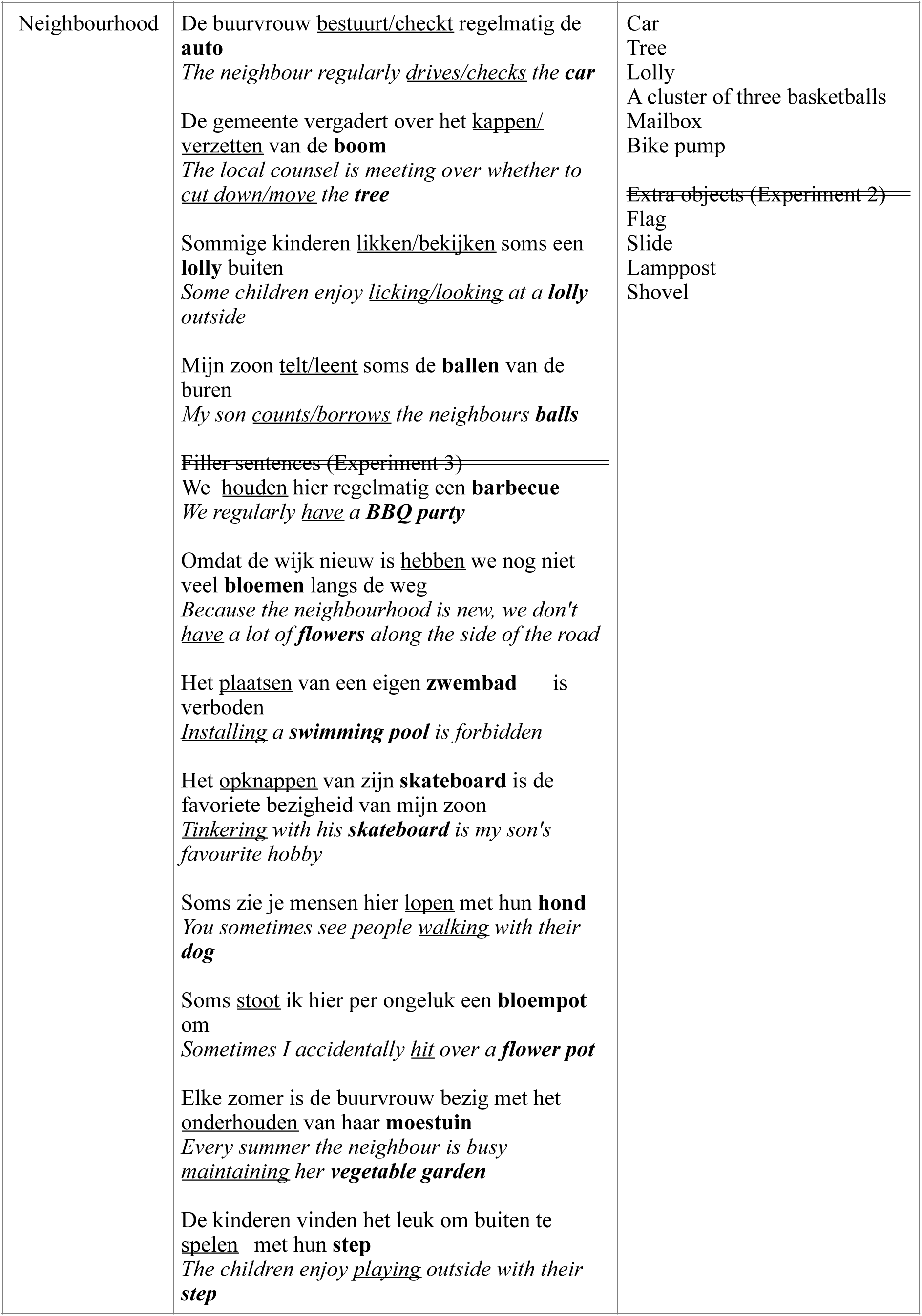

